# Analysis of circulating protein aggregates reveals pathological hallmarks of amyotrophic lateral sclerosis

**DOI:** 10.1101/2020.04.30.070979

**Authors:** Rocco Adiutori, Fabiola Puentes, Michael Bremang, Vittoria Lombardi, Irene Zubiri, Emanuela Leoni, Johan Aarum, Denise Sheer, Simon McArthur, Ian Pike, Andrea Malaspina

## Abstract

Blood-based biomarkers can be informative of brain disorders where protein aggregation play a major role. The proteome of plasma and circulating protein aggregates (CPA) reflect the inflammatory and metabolic state of the organism and can be predictive of system-level and/or organ-specific pathologies. CPA are enriched with heavy chain neurofilaments (NfH), key axonal constituents involved in brain aggregates formation and biomarkers of the fatal neurodegenerative disorder amyotrophic lateral sclerosis (ALS). Here we show that CPA and brain protein aggregates (BPA) from ALS differ in protein composition and appear as a combination of electron-dense large globular and small filamentous formations on transmission electron microscopy. CPA are highly enriched with proteins involved in the proteasome and energy metabolism. Compared to the human proteome, proteins within aggregates show distinct and tissue-dependent chemical features of aggregation propensity. The use of a TMTcalibrator™ proteomics workflow with ALS brain as calibrant reveals 4973 brain-derived low-abundance proteins in CPA, including the products of translation of 24 ALS risk genes. 285 of these (5.7%) are regulated in ALS CPA including FUS (p<0.05). CPA from both ALS and healthy controls affect cell viability when testing endothelial and PC12 neuronal cell lines, while CPA from ALS exert a more toxic effect at lower concentrations. The analysis of resistance to protease enzymes hydrolysis indicates an ALS-specific digestion pattern for NfH using enterokinase. This study reveals how peripheral protein aggregates are significantly enriched with brain proteins which are highly representative of ALS pathology and a potential alternative source of biomarkers and therapeutic targets for this incurable disorder.

**Significance Statement:** Molecular mechanism of neurodegeneration like protein aggregation are important brain-specific alterations which need to be addressed therapeutically. Recently described fluid biomarkers of neurodegenerative disorders provide means for stratification and monitoring of disease progression. Here we show that circulating protein aggregates are easily accessible in blood and reproduce important features of brain pathology for an incurable disorder like amyotrophic lateral sclerosis. They represent a source of biomarkers and of novel therapeutics for ALS.

## Introduction

The lack of biological end-points for stratification of clinically heterogeneous neurodegenerative disorders into more homogeneous and predictable disease phenotypes limits the development of affordable clinical trials (^1^). Investigations into biomarkers for Amyotrophic Lateral Sclerosis (ALS), a fatal neurodegenerative disorder, and for Alzheimer’s disease, have focused on neurofilaments (Nf), tau and beta amyloid proteins based on their propensity to package into complex protein assemblies, including fibrillary formations and aggregates (^2^). As disease propagation in these conditions may be linked to the spread of pathological protein aggregation, the detection in biofluids of aggregate-bound proteins like Nf has been a good strategy for biomarker discovery (^3,4^). Indeed, the change in biofluid levels of Nf in relation to the speed of disease progression and functional impairment, has emerged as a useful tool for the clinical stratification of ALS (^5,6^).

Brain proteins like neurofilaments (Nf) can leak from aggregates-containing neurons and axons undergoing degeneration into extra-cellular fluid, making their way into cerebrospinal fluid (CSF) and blood. In biofluids, proteins like Nf may transiently assemble into circulating protein aggregates (CPA), similarly to what is described for stress granule-like formations in tissues such as brain (^7^). Using ultracentrifugation (UC) and low-complexity binders to extract CPA from plasma of healthy individuals, we have recently shown that CPA are enriched with heavy chain neurofilament (NfH) and not with the light and medium isoforms (NfL, NfM). Overall, protein aggregation is a dynamic phenomenon which is central to the development of neurodegeneration, but it is not well documented in biological fluids (^8^). However, both brain tissue and biological fluids have been reported to show age-dependent increase in protein aggregation (^9^). The age-dependent loss of solubility of proteins in biofluids may relate to the reduction in chaperonal and homeostatic functions that proteins such albumin have (^10^). Age-associated changes in plasma protein composition have recently been investigated in a large cohort of individuals inclusive of a wide age range, leading to the identification of clusters of proteins whose expression is associated with an individual’s biological age and with the health status of different organs (^11,12^).

Our recent mass spectrometry based proteomic study of CPA from individuals with no known neurological conditions has identified mostly cell structural and, to a lesser extent, extra-cellular matrix proteins, while functional analysis has highlighted pathways involved in the inflammatory response and in the phagosome as well as proteins with prion-like behavior (^8^). These biological processes have been reported to be behind the pathology of most neurodegenerative disorders and particularly of ALS (^13–15^). We could therefore speculate that plasma is a carrier of biologically active proteins, which are informative of the physiological and pathological state of organs. Indeed, previous studies have shown that proteins in circulation can influence the regenerative capacity of multiple tissues and organs in mice (^16,17^). If an old mouse is surgically joined to a young mouse in a joint-circulation called parabiosis, many organs, including the brain, can be rejuvenated in the old mouse by some factor passed from the young mouse(^16–18^). It is therefore important to characterize plasma proteins in solution but also compartmentalized in membrane-bound or aggregate-like particles and study their biophysical properties and potential biological effect. This is relevant in neurodegeneration considering that the exchange of a variety of molecular factors between brain and adjacent fluids may be facilitated by an increase in blood brain barrier (BBB) permeability, as reported in ALS and in other neurodegenerative disorders (^19^).

Here we investigate the protein composition as well as the biochemical and biophysical properties of aggregation in plasma from ALS patients and healthy controls (HC). We report a high number of brain-derived proteins within CPA, both in ALS patients and in controls, and the regulation of proteins known to have a pathogenic role in the disease. We also describe an ALS-specific pattern of NfH enterokinase proteolysis in CPA and the biological effect that these formations have in different cell culture models.

## Results

### Study participants

To study CPA protein composition, an initial Mass Spectrometry (MS)-proteomic study was performed on 2 pools of plasma samples (PPS), one containing samples from three fast and three slow progressing ALS individuals and one from six HC individuals (ALS: 5 males (M), 1 female (F); HC: 3M, 3F; age range: ALS: 46.1-78.5; HC 51.8-62.9; Table S1).

For the TMT^®^ proteomic experiment, for protease digestion and for validation by western blot, CPA extracted from plasma samples of distinct and homogeneous ALS and HC cohorts were tested individually (male/female ratio (3:3); age range (ALS: 60.2-68.8; HC: 60.6-68.3; Table S2, S3).

### Extraction of circulating (CPA) and brain protein aggregates (BPA): qualitative analysis by transmission electron microscopy

Using ultracentrifugation (UC), CPA and BPA were extracted from plasma and brain as previously reported (^8^). To verify the efficiency of the aggregate extraction protocols, we have used transmission electron microscopy (TEM) to visualize the aggregate fractions after UC of plasma samples (3 HC and 3 ALS cases) and ALS brains. TEM revealed the presence of (macromolecular) amorphous electron-dense particles of different size, in both CPA and BPA (Figure 1A and B respectively). Despite using the same extraction protocol, in CPA but not in BPA grids, it was possible to appreciate small, round (few nm diameter) and less electron-dense particles close or superimposed to the bigger, globular more electron-dense bodies, likely to represent micelles composed of lipids, detergents or lipoproteins as reported in the literature (^20,21^) (Figure 1 A and B). We could speculate that these particles were made of lipoproteins, such as very low-density lipoproteins (VLDL), low-density lipoproteins (LDL) and high-density lipoproteins (HDL). Lipoproteins were in fact detected by mass spectrometry and biochemical pathways involving these proteins were significantly regulated in the CPA proteomic study (described below). Unlike the large, amorphous, globular appearance of CPA aggregates, some of the electron-dense formations in the BPA micrographs had filamentous and donut-like shapes (Figure 1 B and C), which could be related to the previously reported possible contamination of brain homogenates with ferritin (^22–24^). In CPA grids only, it was possible to see filamentous fragments with a rough surface similar to those seen in BPA (Figure 1D, E and F).

**Figure 1.**
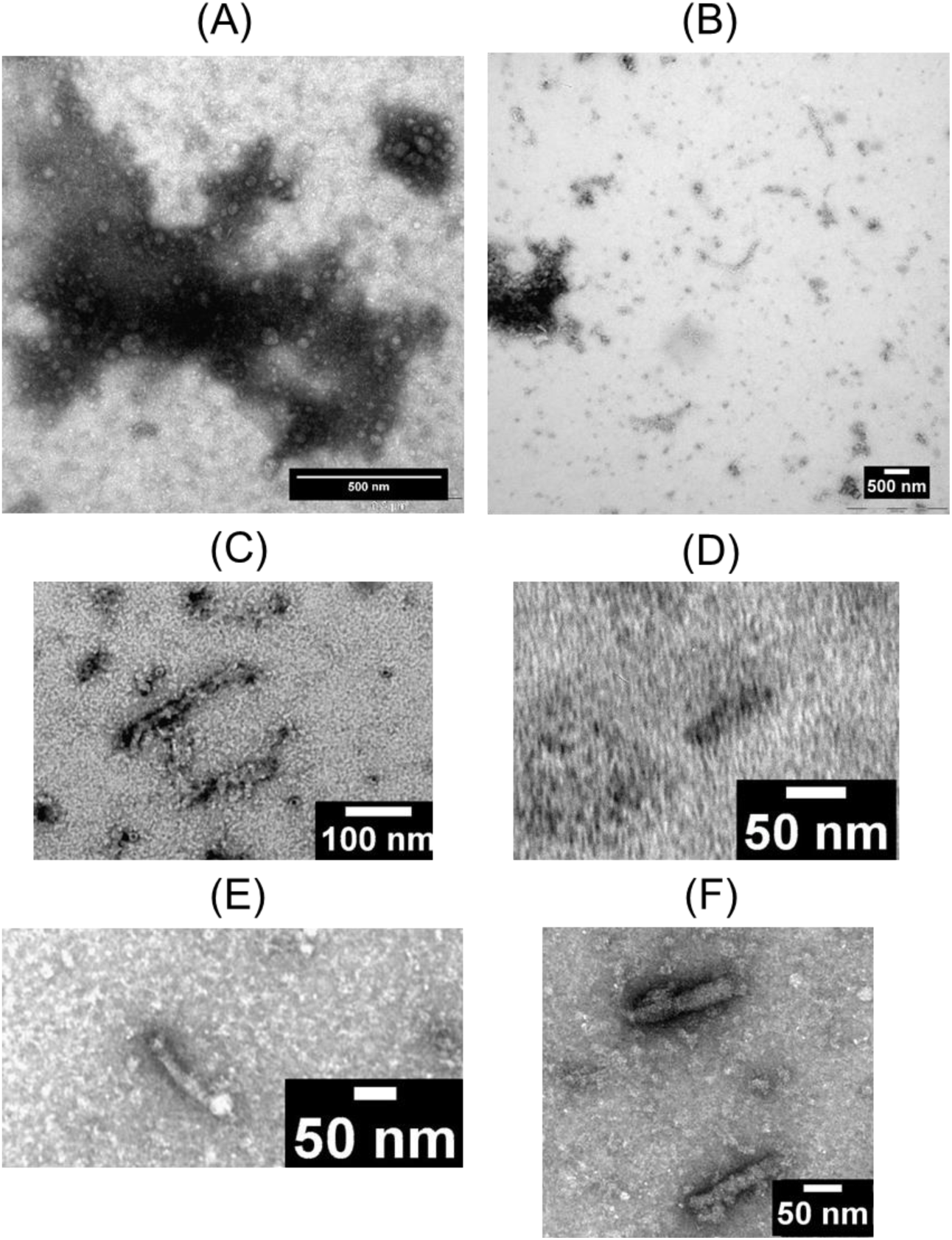
Micrographs of circulating protein aggregates (CPA) and brain protein aggregates (BPA) taken by transmission electron microscopy (TEM) after uranyl acetate (UA) negative staining. (A) grid micrograph after CPA sample loading showing an amorphous globular formation with adjacent and/or superimposed smaller rounded particles (which may be formed of lipoproteins). (B) grid micrograph of BPA showing amorphous electron-dense (left-hand side) as well as short filamentous and small round formations. (C) Details of filamentous and of donut-like particles detected in BPA micrographs. (D, E, F) Micrograph grids of CPA showing 13 to 20 nm thick and 70 to 145 nm long fragments (scale bar on the lower right-hand corner of each micrograph.).

### Circulating protein aggregates (CPA) and brain protein aggregates (BPA) composition: LC-MS/MS proteomics

Liquid Chromatography coupled with Tandem Mass Spectrometry (LC-MS/MS) after in-gel trypsin digestion was used to study protein aggregates enriched from ALS and HC pooled plasma samples (PPS) as well as from ALS brains.

In total, 367 proteins were identified in ALS CPA and 353 in HC CPA (Figure 2A). 264 (57.9% of the total) proteins were expressed in both ALS and HC CPA (here defined as shared), while 103 (22.6%) were found only in ALS (defined as unique ALS) and 89 (19.5%) in HC CPA (defined as unique Controls) (Figure 2A). Functional analysis of these proteome sub-sets was performed using Webgestalt for Kyoto Encyclopaedia of Genes and Genomes (KEGG) pathway enrichment.

**Figure 2.**
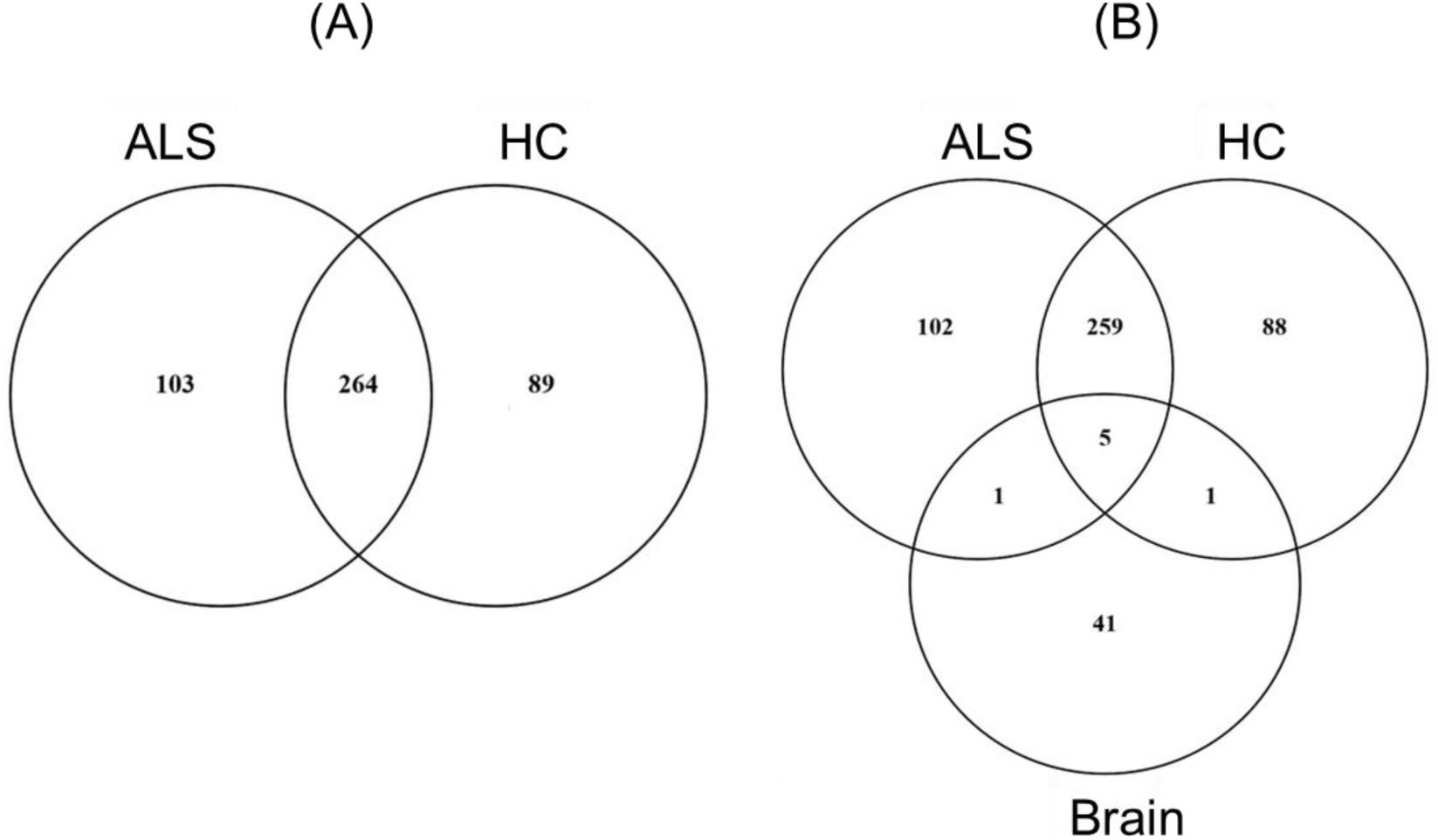
Proteins identified by LC-MS/MS in circulating protein aggregates (CPA) enriched from ALS and HC pooled plasma samples and from aggregates enriched from brain. (A) Venn diagram showing CPA proteins unique to or shared by ALS and HC. (B) Venn diagram showing HC and ALS CPA proteins shared by brain aggregates. Five proteins were expressed in all 3 aggregate groups (actin cytoplasmic 1, tubulin alpha-4A chain isoform 2, clathrin heavy chain 1 isoform 2, collagen alpha-1(VI) and plectin isoform 7), while brain aggregates shared only one protein with ALS and HC CPA (cytoplasmic dynein 1 heavy chain 1 and collagen alpha-2(VI), respectively).

Among the most enriched pathways, the proteasome was the most significantly represented feature in ALS (p=0.028; four genes matched in this category) while the glycolysis/gluconeogenesis pathway (p=0.009; seven genes matched), the pentose phosphate pathway (p=0.003; five genes matched) and the carbon metabolism (p=0.008; eight genes matched) was significantly expressed in HC. Proteins previously linked to ALS like NfH or TDP-43 were not detected.

LC-MS/MS identified 48 protein groups in brain protein aggregates (BPA), including the three neurofilament isoform proteins. There was little overlap between CPA and BPA proteins (Figure 2B), with only five proteins identified in both types of aggregates, one identified in both ALS CPA and BPA and one in both HC CPA and BPA.

#### Aggregation propensity

The brain and plasma aggregate protein lists generated by LC-MS/MS were studied to test the propensity of aggregation in each dataset. The distribution of protein size or molecular weight (MW), isoelectric point (pI) and hydrophobicity, by the GRAVY index, were analysed in each proteome dataset and the whole human proteome was used as reference (^25^). BPA had a significantly higher MW (p<0.0001) compared to the other datasets (ALS, HC, shared respectively and Human proteome, Figure 3A), while pI was significantly lower in all CPA datasets compared to the Human proteome (p <0.0001; Figure 3B). Despite minimal overlap in protein composition between CPA and BPA (Figure 2B), aggregation propensity in the two aggregate types and in the human proteome was similar when measured by GRAVY index, which takes into account the average hydropathy of a peptide according to its aminoacidic composition (Figure 3C).

**Figure 3.**
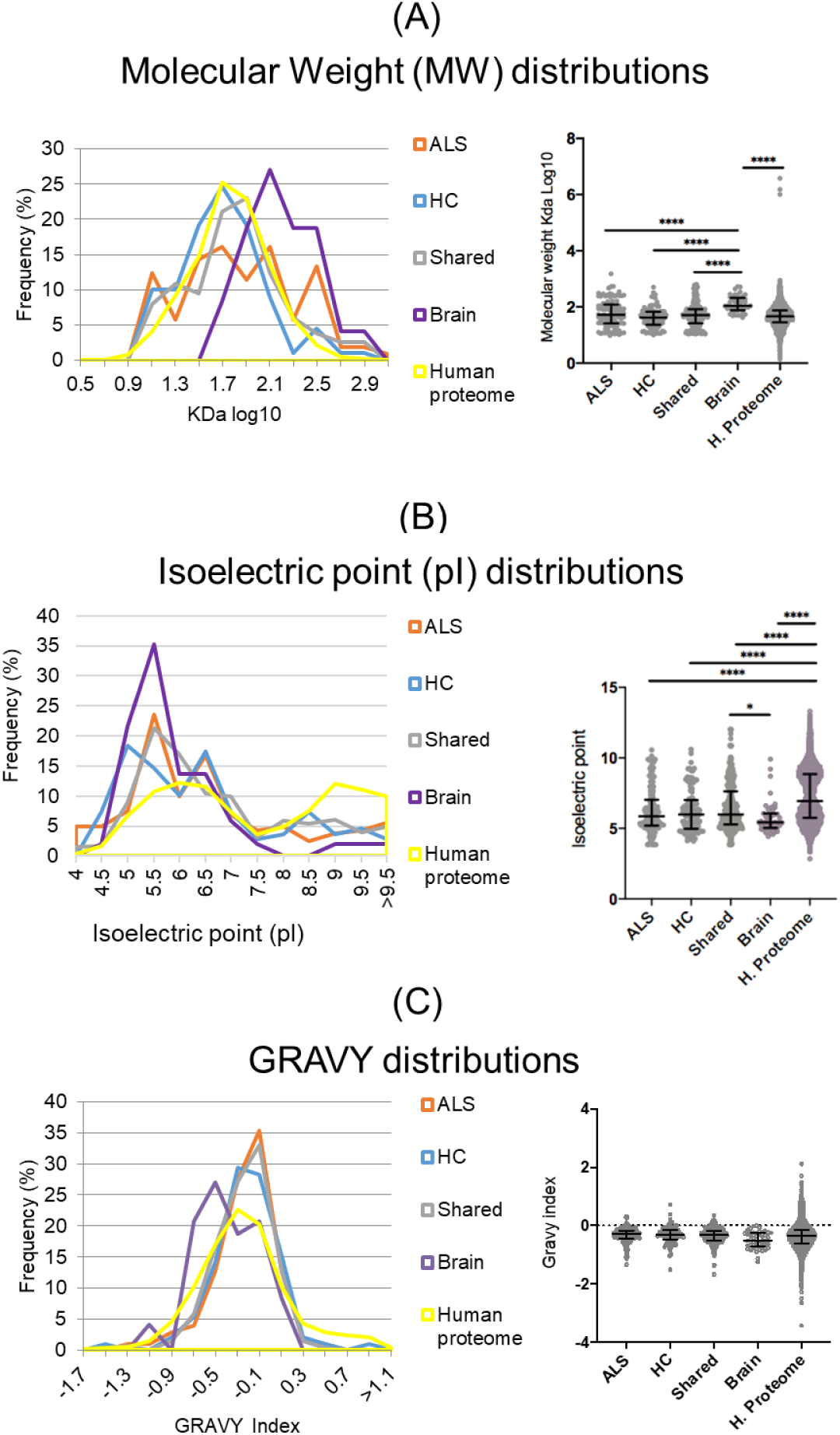
Aggregation propensity of proteins in circulating and brain protein aggregates from ALS and HC compared to the Human proteome. Molecular weight (MW) (A), isoelectric point (pI) (B) and hydrophobicity (GRAVY index) (C), known to affect aggregation propensity, are compared across proteins found to be expressed only in ALS and HC CPA (ALS and HC respectively), proteins shared between ALS and HC CPA datasets (Shared), proteins within brain aggregates (Brain) and in the entire human proteome. The distribution plots show the dispersion of the samples with relative frequency, while the violin plots show median and interquartile ranges. Statistical analysis was performed on ALS, HC, Shared and Human proteome with one-way ANOVA, Kruskal-Wallis test with Dunn’s multiple comparison as post-test for group analysis with * expressing the level of significance (*: p = 0.0251; ****: p < 0.0001).

### CPA protease digestion and NfH resistance

Resistance to protease digestion has been described as a key feature of altered protein behavior in conditions like prion disease (^26^). We have previously shown that neurofilament heavy chain (NfH) is found in blood CPA (^8,27^). As NfH is constitutively expressed in protein aggregates from ALS brain and has been linked to the pathogenesis of the disease, we have looked at NfH protease resistance in CPA from ALS and compared its digestion profile to HC (Figure 4). CPA enriched from the ALS and HC plasma samples were treated with trypsin, chymotrypsin, enterokinase and calpain. Both CPA NfH digested and undigested profiles were analysed.

**Figure 4.**
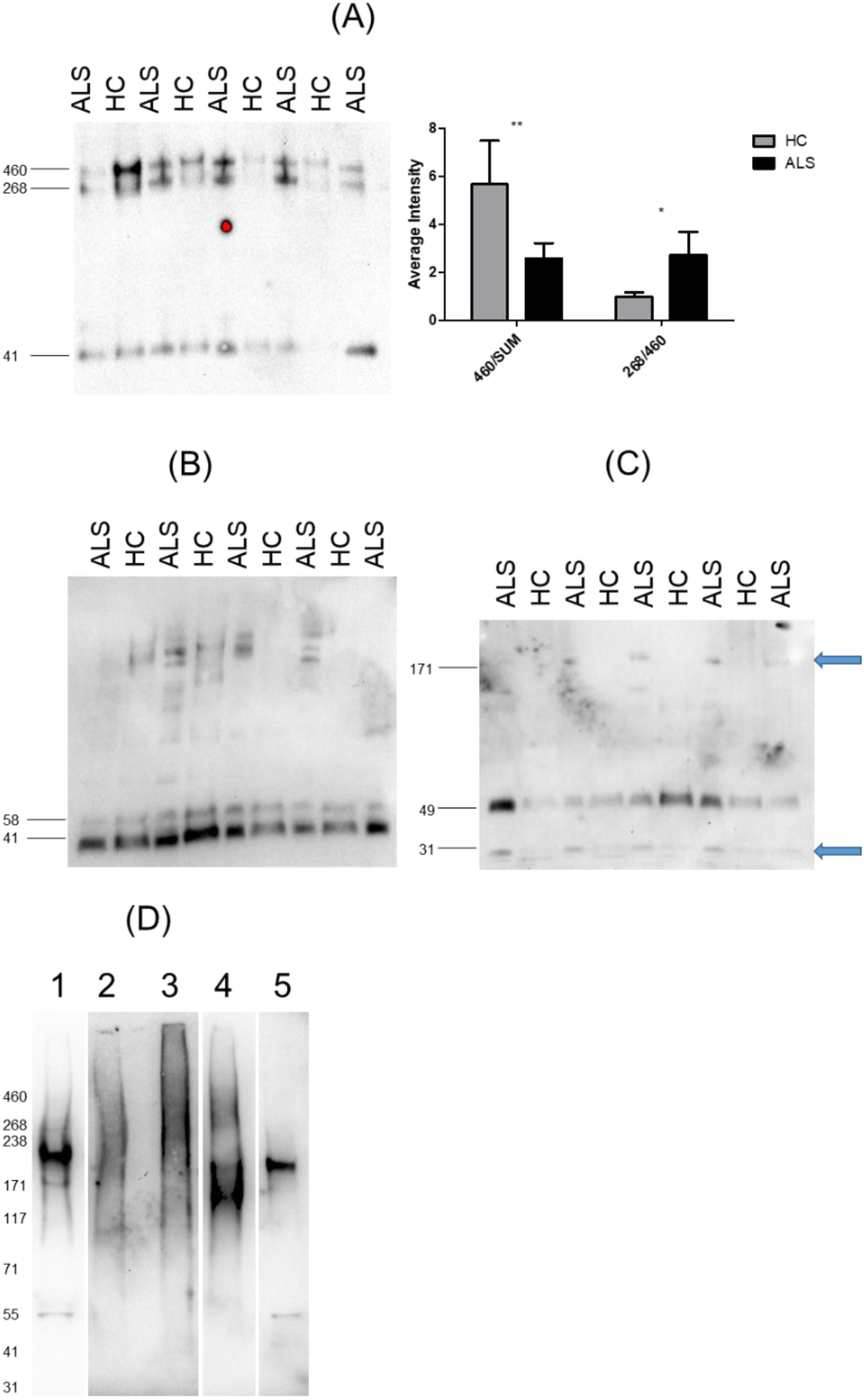
Westerrn blot analysis of neurofilament heavy chain (NfH) within circulating protein aggregates (CPA) after proteases digestion. Undigested CPA (A) show NfH bands at 460, 268 and 41 KDa (268 KDa is NfH expected molecular weight). The ratio between the 460 KDa band and the the sum of all NfH band intensities (SUM, 460/SUM) is higher in HC (p= 0.032), while the ratio between the 268 and 460 bands (268/460) is higher in ALS (p= 0.018). Calpain digestion (B) shows 58 and 41 KDa bands in all samples with no difference in expression. The enterokinase digestion profile of NfH (C) shows a 49 KDa band uniformly expressed across samples and additional 171 and 31 KDa bands only in ALS patients (blue arrows). Undigested NfH in ALS brain protein aggregates (BPA; D, lane 1) and after digestion with chymotrypsin (lane 2), enterokinase (lane 3), calpain (lane 4) and brain lysate lane 5. To maximise band visualization, time exposure was for lane 1 at 10.1 seconds, lane 4 and 5 at 58.4 seconds and lane 2 and 3 at 278.8 seconds.

Western blot analysis of NfH in plasma CPA before digestion detected three bands at 460, 268 and 41 KDa as previously reported (Figure 4A) (^8^). The sum of all NfH band intensities (SUM) was higher but not statistically significant in the ALS group compared to HC. The ratio between the intensity of the 460 KDa band and the NfH SUM intensity (460/SUM) was higher in HC, while the ratio between the intensities of the 268 and 460 bands (268/460) was significant higher in ALS (Figure 4A).

Treatment with trypsin or chymotrypsin showed an almost complete digestion of NfH in both ALS and HC, with the exception of a residual 41 KDa band present in a minority of samples (data not shown). After calpain digestion, western blot analysis showed a different pattern of immunoreactivity for each CPA sample with the exception of 58 and 41 KDa bands evenly detected in all samples (Figure 4B). Enterokinase digestion resulted in a 49 KDa band in all samples with equal expression in ALS and HC (Figure 4C). All ALS samples showed bands at 171 and 31 KDa not seen in HC samples (Figure 4C).

NfH in BPA showed low or no resistance to digestion with all three enzymes (Figure 4D). Chymotrypsin and enterokinase digestions (Figure 4D, lane 2 and 3 respectively) generated no distinct bands but a faint smear at higher molecular weight than the NfH bands detected in undigested brain and brain lysate (Figure 4D, lane 1 and 5 respectively). Calpain digestion showed two faint bands at about 171 KDa (Figure 4B, lane 4).

### TMTcalibrator™: brain-derived proteins in CPA

The observed lack of similarity in the composition of brain and plasma aggregates may relate to the limits of proteomic techniques, whereby low abundance proteins, such as the brain-derived ones, may be masked by those with high abundance and detection confounded by the presence of post-translational modifications. To address these shortfalls and gather more information on the potential enrichment of brain-derived proteins in circulating aggregates, we have undertaken further proteomics using a TMTcalibrator™ workflow, where brain lysate was used to enhance detection of proteins in CPA (^28–30^).

4973 brain-derived proteins were identified, including the three neurofilaments (Nf) protein isoforms (Nf Light (NfL), Nf Medium (NfM) and Nf Heavy (NfH)). Nf were found at a relatively higher level in ALS compare to HC samples (log2-fold change ALS/HC (logFC) = 0.093, 0.181 and 0.298, respectively) but none was significantly regulated (p= 0.40, 0.16 and 0.06 respectively). Of the 4973 brain-derived proteins, 285 proteins (5.7%) showed a statistically significant regulation (p < 0.05). 158 were more expressed in HC (logFC < 0) with an average fold-change (FC) of −0.667, while 127 in ALS (logFC > 0) with an average FC of 0.703. The protein list obtained was matched with an ALS gene list obtained from the MalaCards database (^31^), an integrated repository of human diseases and their annotations. 24 ALS proteins were identified including Fused in Sarcoma RNA-binding protein (FUS) which was found to be significantly regulated in ALS (p= 0.00696) (Table 1).

**Table 1.** ALS risk genes included in the list of proteins identified by the TMTcalibrator™ workflow in CPA from ALS and HC and reported in the gene classifiers MalaCards Human Disease Database (https://www.malacards.org/card/amyotrophic_lateral_sclerosis_1#RelatedGenes-table) (^31^). Among a total of 38 ALS elite genes reported in the MalaCards Human Disease Database (those more likely to cause the disease), 24 were detected in the list of proteins generated by the TMTcalibrator™ experiment. A: the gene symbol used to represent a gene B: Uniprot database protein identifier C: protein full name recommended by Uniprot D: number of peptide sequences unique to a protein group E: relative quantification with value expressed as log2(ALS/HC) intensities F: statistical significance for differential regulation between ALS and HC experimental groups

| <sup>A</sup>Gene | <sup>B</sup>Uniprot ID | <sup>C</sup>Protein name | <sup>D</sup>Unique peptides | <sup>E</sup>logFC | <sup>F</sup>p-value |
| --- | --- | --- | --- | --- | --- |
| **FUS** | P35637-2 | Isoform Short of RNA-binding protein FUS | 5 | 0.564 | 6.96e<sup>-03</sup> |
| **NEFH** | P12036 | Neurofilament heavy polypeptide | 27 | 0.298 | 6.36e<sup>-02</sup> |
| **OPTN** | Q96CV9 | Optineurin | 11 | -0.243 | 9.58e<sup>-02</sup> |
| **UNC13A** | Q9UPW8 | Protein unc-13 homolog A | 11 | -0.326 | 9.75e<sup>-02</sup> |
| **PON2** | Q15165-3 | Isoform 3 of Serum paraoxonase/arylesterase 2 | 1 | 0.292 | 1.46e<sup>-01</sup> |
| **ANG** | P03950 | Angiogenin | 1 | -0.595 | 1.67e<sup>-01</sup> |
| **CHMP2B** | Q9UQN3 | Charged multivesicular body protein 2b | 1 | 0.492 | 1.90e<sup>-01</sup> |
| **VCP** | P55072 | Transitional endoplasmic reticulum ATPase | 65 | -0.514 | 2.16e<sup>-01</sup> |
| **ATXN2** | Q99700-2 | Isoform 2 of Ataxin-2 | 2 | -0.445 | 2.51e<sup>-01</sup> |
| **ANXA11** | P50995-2 | Isoform 2 of Annexin A11 | 16 | 0.150 | 3.01e<sup>-01</sup> |
| **SOD1** | P00441 | Superoxide dismutase [Cu-Zn] | 8 | 0.176 | 3.39e<sup>-01</sup> |
| **ERBB4** | Q15303-4 | Isoform JM-B CYT-2 of Receptor tyrosine-protein kinase erbB-4 | 1 | -0.284 | 3.70e<sup>-01</sup> |
| **TARDBP** | Q13148 | TAR DNA-binding protein 43 | 2 | 0.180 | 3.85e<sup>-01</sup> |
| **SQSTM1** | Q13501 | Sequestosome-1 | 2 | -0.280 | 4.00e<sup>-01</sup> |
| **MATR3** | P43243 | Matrin-3 | 16 | 0.127 | 4.03e<sup>-01</sup> |
| **PFN1** | P07737 | Profilin-1 | 14 | 0.136 | 4.08e<sup>-01</sup> |
| **VAPB** | O95292 | Vesicle-associated membrane protein-associated protein B/C | 10 | -0.094 | 4.31e<sup>-01</sup> |
| **EPHA4** | P54764 | Ephrin type-A receptor 4 | 13 | -0.082 | 5.37e<sup>-01</sup> |
| **PON1** | P27169 | Serum paraoxonase/arylesterase 1 | 14 | 0.123 | 6.07e<sup>-01</sup> |
| **TAF15** | Q92804-2 | Isoform Short of TATA-binding protein-associated factor 2N | 3 | 0.067 | 6.99e<sup>-01</sup> |
| **UBQLN2** | Q9UHD9 | Ubiquilin-2 | 4 | -0.065 | 6.99e<sup>-01</sup> |
| **HNRNPA1** | P09651-3 | Isoform 2 of Heterogeneous nuclear ribonucleoprotein A1 | 8 | -0.066 | 7.50e<sup>-01</sup> |
| **DCTN1** | Q14203-6 | Isoform 6 of Dynactin subunit 1 | 2 | 0.020 | 9.34e<sup>-01</sup> |
| **TBK1** | Q9UHD2 | Serine/threonine-protein kinase TBK1 | 5 | -0.004 | 9.86e<sup>-01</sup> |

Principal component analysis (PCA) of the proteomic data after correction for the TMT^®^ batch-effect identified the sample class (ALS or HC) as the strongest component of the variance in the data matrix (41.17% of the total variance, Figure 5A). The marked separation between ALS and HC in CPA brain-derived proteins identified by PCA was also observed when using clustering of these regulated proteins (Figure 5B). Considering the CPA enrichment of brain proteins (Figure 5C), we speculate that circulating protein assemblies may represent a good source of biomarkers for ALS.

**Figure 5.**
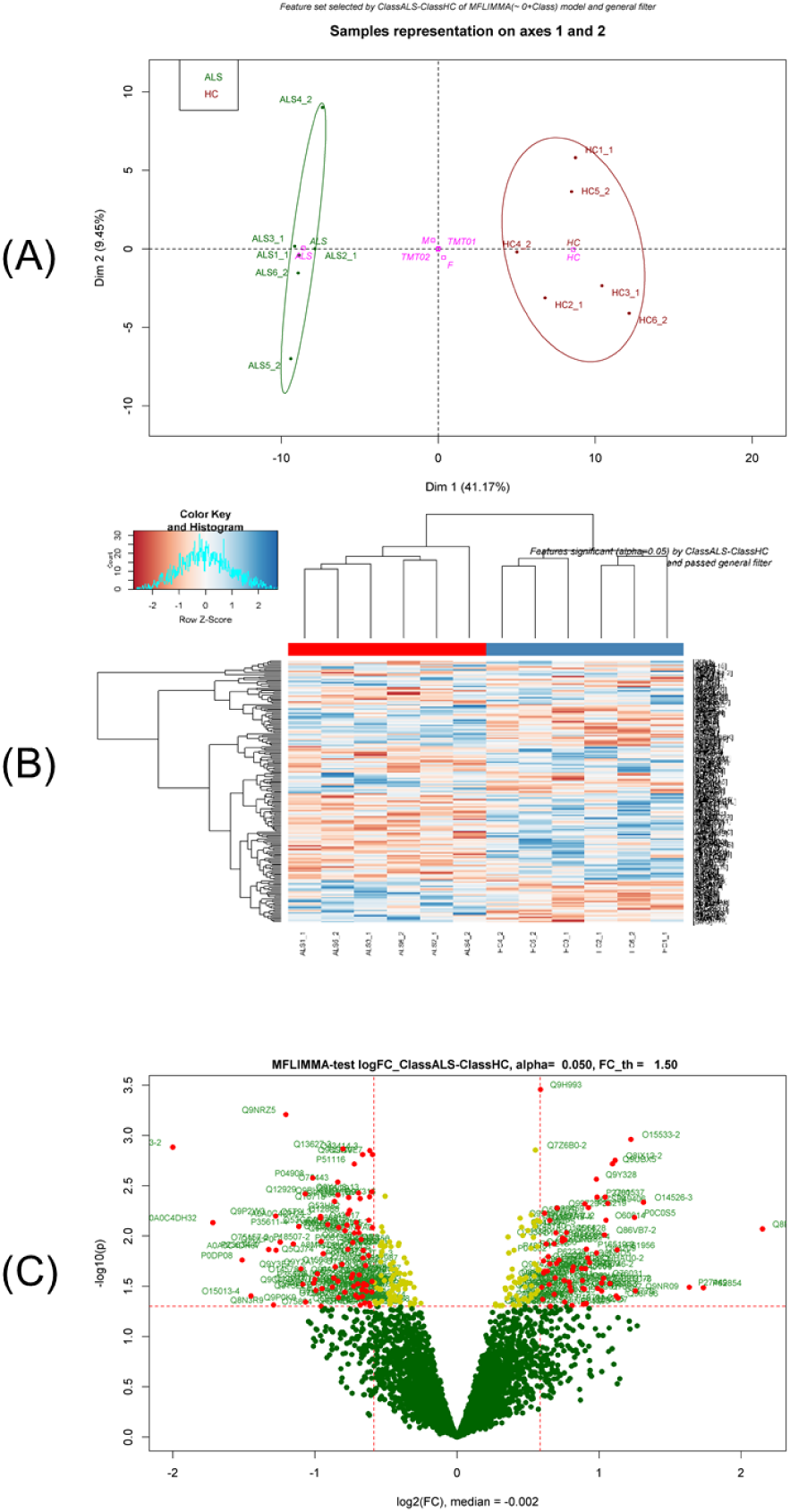
TMTcalibrator™ proteomic analysis. (A) Principal component analysis (PCA) showing a separation between the ALS and HC experimental groups regulated features at protein level. Dimension 1 or the variance between the two experimental groups (ALS and HC) is 41.17% of the entire variance; dimension 2 or variance between 10plexes (TMT01 and TMT02) is 9.45% of the entire variance. (B) Heatmap showing the distribution of the regulated features and their clustering. The regulated features are distributed vertically, reported as Uniprot IDs on the right-hand side and relative clustering on the left-hand side. Analytical samples are distributed horizontally, with sample names at the bottom and relative clustering at the top of the heatmaps. The color key histogram at the top left side shows the distribution of the features and the heatmap color coding. (C) The volcano plot shows the distribution of the proteins identified by TMT proteomic study according to their fold change (FC) expressed as log2 (fold change ALS/HC) (logFC) in the x axis and according to p-value expressed as –log10 (p-value) in the y axis. Protein groups were considered regulated if p-value < 0.05 and logFC < −0.58 or > 0.58. Red dots are regulated features, yellow dots are features with a significant p-value (p < 0.05) and logFC between −0.58 and 0.58 while green dots are not regulated protein groups (p > 0,05). Uniprot IDs are reported beside the dots with significant p-value.

#### Functional analysis

We applied different approaches to assess the relevance of different biological terms, based on their over-representation in the subset of regulated proteins (^28,32^). Among the biochemical pathways identified, 69 showed a p-value < 0.05. Within the top ten regulated pathways, five were involved in metabolism of lipoproteins (Table S4). The significant representation of these pathways in our brain/CPA proteomes reflects known changes of the ALS pathophysiology, where metabolism is thought to be switched from sugars and carbohydrates to lipids use (^33–35^). To identify potential protein candidates for biomarkers analysis, we looked at highly regulated protein (unique peptides ≥ 2, LogFC < −0.693 or > 0.693, p-value < 0.05; 48 proteins in total) within the 69 pathways highlighted by FAT. Four biochemical pathways seemed to represent the most consistent changes at a proteomic level: metabolism of carbohydrates (p= 0.0099), glycosaminoglycans (GAGs) metabolism, lysosome (p= 0.0015), synthesis of phosphatidic acid (p= 0.0184) and wnt signalling pathway (p= 0.0337). It has been shown that GAGs are involved in protein aggregation and prion diffusion (^36–44^). Lysosome activity changes were described in ALS caused by the C9orf72 gene repeat expansions (^45–49^), while synthesis of PA is linked to all types of phospholipids, including phosphatidylcholine and phosphatidylethanolamine that have been linked to ALS and prion disease pathogenesis (^50,51^).

#### Aggregation propensity of the brain-derived proteins in CPAs

We evaluated aggregation propensity of the CPA proteins identified by TMTcalibrator™ compared to the Human proteome and BPA, using the reported physicochemical parameters. The analysis was performed on the entire TMT^®^ proteome dataset and regulated proteins only. Among the latter subset of proteins, two groups were considered based on the differential expression: log(ALS/HC)>0 and log(ALS/HC)<0. There was no statistically significant difference among these three datasets for the parameters under investigation. However, the entire TMT and BPA proteome datasets showed statistically significant higher MW and lower pI compared to the Human proteome (P <0.0001), while for GRAVY index, the BPA values were lower than TMT and Human proteome (p= 0.0348 and 0.0224, respectively; Figure 6).

**Figure 6.**
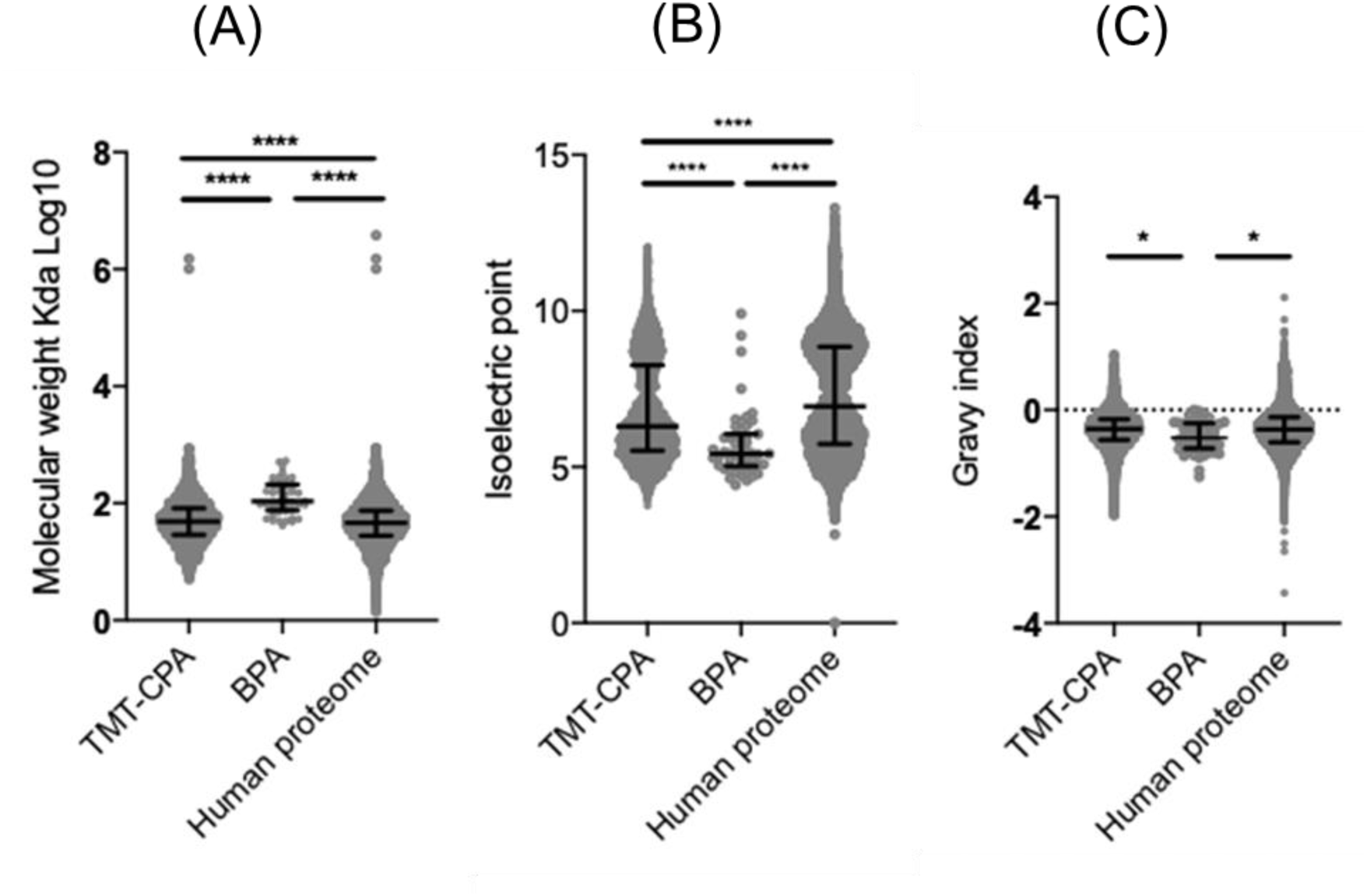
Comparison of aggregation propensity parameters of CPA proteins obtained through the TMT proteomic workflow (TMT-CPA), BPA proteins by LC-MS/MS and the Human proteome. Molecular weight (MW) (A), isoelectric point (pI) (B) and hydrophobicity (GRAVY index) (C) were compared across TMT-CPA, BPA and Human proteome datasets. Statistical analysis was performed with one-way ANOVA, Kruskal-Wallis test with Dunn’s multiple comparison as post-test. The violin plots show median and interquartile ranges across the three datasets. The TMT-CPA dataset differ significantly compared to the Human proteome and BPA for MW and pI, while for hydrophobicity (GRAVY index), there is only a less significant difference with the BPA dataset. **** p < 0.0001. (A and B); * p = 0.0348, TMT-CPA versus BPA (C); * p = 0.0224, BPA versus Human proteome (C).

#### Proteomic results validation by immunoassays

Six proteins belonging to ALS-relevant molecular pathways were tested by western blot: Glypican-4 (GPC4), Fibromodulin (FMOD), Biglycan (BGN), Cation-dependent mannose-6-phosphate receptor (M6PR), Endophilin-B2 (SH3GLB2) and Protein DJ-1 (PARK7) using CPA extracted from ALS patients and HC, with brain lysate as reference. SH3GLB2 showed the same trend of ALS vs HC protein regulation identified in the TMTcalibrator™ experiment, with a similar level of regulation (logFC= 0.34 in TMTcalibrator™ and log2(ALS/HC)= 0.437) in western blot analysis (Figure 7). The remaining proteins showed a different trend of regulation compared to that obtained in the proteomic analysis (data not shown). SH3GLB2 (as well as M6PR) were detected at a higher MW in CPA compared to brain lysate, possibly in keeping with a different PTM profile in tissues as opposed to fluids which may affect the state of aggregation and protease digestion.

**Figure 7.**
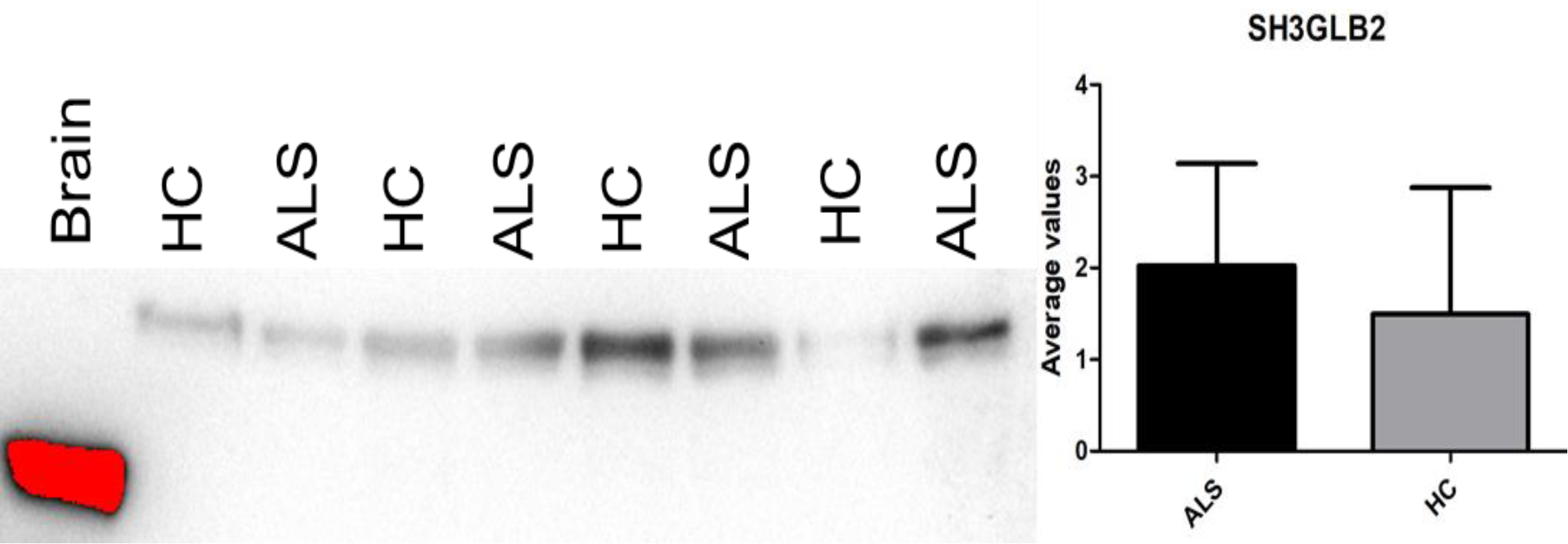
Western blot analysis of Endophilin-B2 (SH3GLB2) in plasma CPA from ALS patients and healthy controls. Samples were normalized to HC6 density and the average values with relative standard deviation for the ALS (n=4) and Control (n=4) groups were plotted onto the chart. A brain lysate sample is also included (1st lane, red band, indicating signal saturation) which showed an endophilin-B2 band at a lower molecular weight than the bands detected in CPA. Immunodetection confirmed the SH3GLB2 higher level of expression in the ALS CPA compared to control (logFC= 0.34), but without being statistically significant (p= 0.57).

### Cell survival assays

To test the biological effects of CPA on living cells, human brain microvascular endothelial cells (hCMEC/D3) modelling human blood-brain barrier (BBB) and PC12 neuron-like cells lines were treated with CPA from ALS patients and HC and cell viability measured (Figure 8). Total IgG were extracted from the same blood samples CPA were separated from and used to treat the same cell lines. CPA re-suspended in PBS were administered at defined concentration to hCMEC/D3, while CPA were pre-treated with 8M urea before testing PC12 viability. Dissolution of aggregates by urea for PC12 cells was undertaken to evaluate the effect of CPA-containing proteins rather than the effect of their aggregated state. PC12 cell viability decreased at increasing concentration of CPA, (to 75% at 0.5 µg/ml; Figure 8B), while endothelial cells showed the opposite trend, with the maximum effect on cell viability at the lowest concentration (0.05 µg/ml) and no effect at the highest concentration (1 µg/ml; Figure 8A). ALS CPA exerted higher toxicity (lower cell viability) at lower concentration (0.05 µg/ml) with endothelial cells and with PC12 cells (0.5 µg/ml) compared to HC CPA (Figure 8). IgG extracted from the same ALS and HC plasma samples as CPA, showed reduction of endothelial cell viability to 80% and to between 70 and 85% in PC12 cells at 1.5 µg/ml, with no significant difference between ALS and HC (Figure 8).

**Figure 8.**
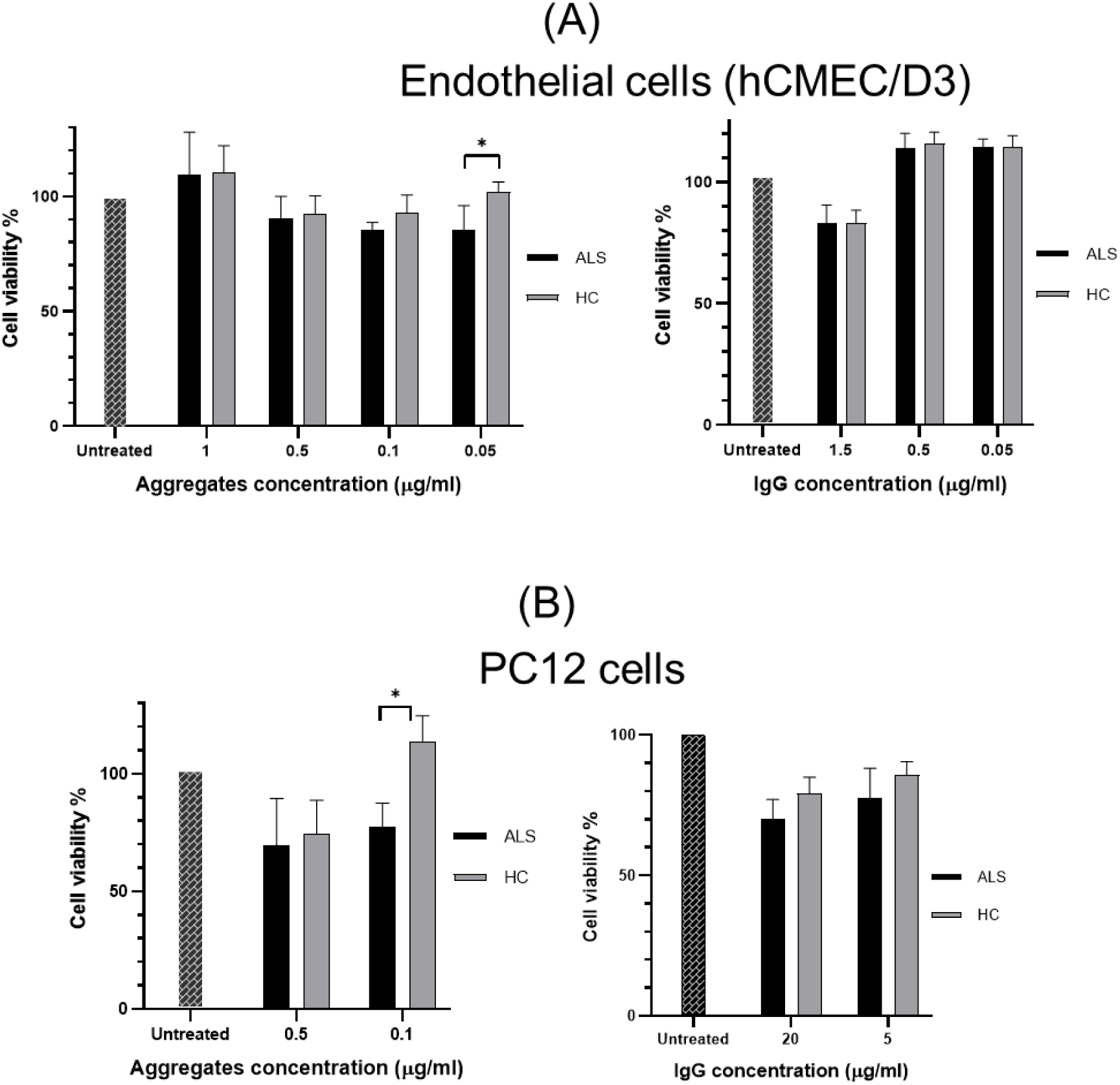
Cell viability after treatment with circulating protein aggregates (CPA), proteins solubilized from CPA and immunoglobulins extracted from the same plasma samples. The figure shows the percentage of endothelial (hCMEC/D3) and PC12 living cells (A and B) after treatment with different concentrations of CPA and IgG from ALS and HC. Cells treated with ALS CPA showed a statistically significant lower cell viability compared to HC CPA treated cells at 0,05 µg/ml (p= 0,031; endothelial cells, A) and at 0,1 µg/ml (p= 0,029; PC12 cells, B). IgG had minor effect on all cell type viability with no difference between ALS and HC. CPA proteins were solubilized with 8M urea before PC12 cells treatment. Significance was tested by two-way ANOVA and Tukey HSD test.

## Discussion

Here we investigate protein aggregation in biofluids with reference to the same process in brain tissue. In ALS, an incurable neurodegenerative disorder characterized by a rapid spread of neuroaxonal loss, we show that circulating protein aggregates (CPA) become enriched with proteins involved in the proteasome, a clearance mechanism of defective proteins implicated in the disease pathogenesis (^52,53^). Using a pioneering proteomic approach to study the interface between fluid and brain tissue, the TMTcalibrator™, we show for the first time, that CPA contain just below 5,000 brain proteins of which 285 are regulated in ALS, including products of translation of ALS risk genes such as FUS and SOD1 (Table 1). As for brain aggregates, the CPA protein mix shows features of increased propensity to aggregate compared to the human proteome as a whole, while CPA from ALS plasma exert a toxic effect at a low concentration on both endothelial and neuronal cell lines which exceeds the effect of CPA from HC on the same cell types.

The TMTcalibrator™ workflow employed in this study has the added value of being able to analyze tissues and fluids in the same round of experiments, enhancing the detection of fluid proteins that are also expressed in brain tissue, most of them usually undetectable using standard proteomic approaches (^8^). Within the large number of brain-derived proteins in CPA, 285 show a significant level of regulation (p < 0.05) in ALS compared to controls (Figure 5). The detection of a large subset of low-abundance brain proteins in circulating molecular assemblies has significant conceptual implications also from a methodological standpoint. Immunodetection, for example, would not normally reliably detect in the same experiment such a variety of proteins and, as recently shown for neurofilaments, it can suffer from competition from naturally occurring antibodies which cause epitope sequestration in aggregates and immunocomplexes (^5,27^). Unlike, TMTcalibrator™ enhanced detection based on an internal tissue calibrant, standard MS-based proteomics would, in turn, lack the sensitivity to discriminate low concentration proteins against those that are more abundant. Whilst more work needs to be done to test how the expression of these neuronal protein change in biofluids under pathological conditions affecting the brain, the identification in blood of such a wide range of disease-relevant neuronal constituents provides, without doubt, the basis for a novel strategy for biomarker discovery in ALS.

The limitation of our study is the inability to determine whether the target proteins are mostly in an aggregated or soluble state. The method of aggregates separation employed in our study does not allow investigation by proteomics of the non-aggregated proteome (supernatant) due to technical limitations in the CPA extraction. To circumvent this problem, we have previously used low complexity binders for aggregate separation in biofluid, only to observe that the resulting aggregate fraction had a substantially different composition to the one obtained by ultracentrifugation, the method of choice in the work presented here (^8^). Our recent use of two separate proteomic workflows to study the immunological response and the plasma/brain proteome in phenotypic variants of ALS allows further interpretation of our data (^28^). In this study, we have used the same brain-enhanced TMT proteomics platform to test neuronal-derived proteins expression in whole plasma, and not in the plasma aggregate-fraction only. We have identified nine ALS elite genes that are among the 24 identified in our CPA study (including Profilin-1 and the Isoform 2 of Heterogeneous nuclear ribonucleoprotein A1). This may indicate that aggregates in blood represent a state of enrichment of brain and disease-specific proteins, certainly in the case of ALS, compared to proteins in a fluid state. However, low abundance neuronal proteins would ultimately make up a small component of these macromolecular formations and require sensitive proteomic techniques for their detection.

Unlike other studies of the whole plasma proteome in ALS, here we show that the protein composition of plasma CPA from ALS individuals reproduces most of the extensively described pathological hallmarks of ALS, including pathways involved in protein degradation through the proteasome and energy metabolism (^52–55^). The same approach applied to the TMT-proteomic datasets reveals regulated features also known to be involved in ALS pathology and in protein aggregates homeostasis including lysosome as well as lipoprotein and glycosaminoglycan metabolisms (^33,34,45,48,49,52–55^) (Table S4). All disease-related alterations in proteasome activity in ALS have so far been shown in such detail only in brain, spinal cord and in neuronal cell lines, but not systemically or more specifically in blood, as in our study (^52,53^). Our data ultimately points to the protein degradation molecular pathway as a critical alteration in ALS which is detectable as a systemic response. To our knowledge, there is no previous description of this phenomenon in the literature.

The analysis of the chemical properties of plasma and brain aggregates provides further insight into protein behavior in different molecular environments and the relation with a disease state like ALS. Using well-established physicochemical parameters of the propensity of proteins to aggregate, we have compared the proteomes of the aggregates under investigation to look at differences based on tissue/fluid origin or on the presence of a pathological state like ALS. When molecular weight and isoelectric point are taken into account, our data show that the main contributor to the differences in chemical properties observed across aggregate types is not the disease state but rather the tissue of origin (Figure 3 and 6). Refence to the whole human proteome also confirms that proteins falling into an aggregate state, regardless of the tissue of origin or presence of a pathological state, make up defined sub-proteomes that can be distinguished from the whole set of human proteins.

The presence of NfH within CPA and the different amount of specific immunoreactive species (Figure 4A) is an important finding from a biomarkers perspective. Moreover, when hydrolytic properties are analysed separately in ALS and HC CPA for NfH, clear differences are identified, especially after enterokinase digestion including the presence of specific fragments at 171 KDa and at 31 KDa only in ALS samples (Figure 4C). While a trend for NfH proteolytic fragments over-expression in ALS CPA compared to HC is visible in our experimental data, our study is not powered with enough samples to establish this observation along what already previously reported (^5,56^). An extension of this preliminary finding using a larger number of samples will be needed to compare our observation to previously reported data on the same neurofilament isoform enterokinase digestion pattern in a different experimental context (^57^).

We have tested different cell lines using change in cell viability upon treatment with ALS and HC CPA as a readout of a potential biological effect of circulating aggregates (Figure 8). Given that immunoglobulins make up a sizeable share of plasma proteins and that plasma has been shown to have a biological effect on tissues, we have treated the same cell lines with immunoglobulins extracted from the same blood samples CPA were recovered from. Whilst immunoglobulins seem not to significantly interfere with cell function at different concentration, there is a detectable toxic effect on PC12 neurons and on hCMEC/D3 endothelial cell cultures upon treatment with CPA, and a clear ALS-specific change in cell viability when CPA are administered at relatively low concentrations (Figure 8). Therefore, CPA may well be one of the proteinaceous component in blood that is ultimately responsible for the BBB damage that has been reported in ALS and in other neurodegenerative conditions (^19^). Based on the experiments described in this paper, it is not possible to explain how endothelial cell damage comes into play upon exposure to CPA, and whether the ALS-specific effect relates to a particular composition and/or conformation of aggregates which may be concentration-dependent. The use of TEM to confirm the efficiency of aggregates separation confirms the presence of particles of both globular and filamentous appearance, similar to those observed in BPA (Figure 1). Taking into account the potential pitfalls of aggregates extraction using ultracentrifugation, we believe that further investigation by TEM are required to understand whether the smaller and filamentous formations are more represented in samples from ALS or from specific ALS phenotypes, given the possible higher level of toxicity when proteins assemble as oligomeric formations. Immuno-gold TEM with appropriate negative controls could also be employed to evaluate CPA relative composition in specific protein types.

Our data support a new concept of biomarker discovery for neurodegenerative disorders based on the enrichment of brain proteins in circulating aggregates and on the change in expression and biophysical properties that most disease-relevant proteins may have under pathological conditions. To date, there is no evidence in the literature of any investigation into non-membrane bound particles in blood stream of ALS patients and animal models. This is important considering our data suggesting that aggregates extracted from ALS plasma affect endothelial and neuronal cell viability when administered to cell culture. Further investigation on the nature of these particles will be required to confirm and strengthen this finding, including a more extensive comparison with brain aggregates and analysis of the biochemical characteristics in a larger subset of individuals, using less labor-intensive but equally effective methods of CPA extraction.

## Materials and Methods

### Patients and biological samples

Samples were collected from individuals with a diagnosis of amyotrophic lateral sclerosis (ALS) according to established criteria (El Escorial Criteria (^58^) and healthy controls (HC), enrolled in the ALS biomarkers study (REC n. 09/H0703/27). Participants had no known neurological comorbidities, nor were they affected by systemic or organ-specific autoimmune disorders (Table S1-3). Blood was drawn by venipuncture in EDTA tubes, processed within 2 hours by spinning at 3500 rpm for 10 minutes at 20 °C and stored at −80 °C.

Pre-central Gyrus brain tissue samples from two individuals affected by ALS (Brain1 and Brain2) obtained from The Netherlands Brain Bank (Netherlands Institute for Neuroscience, Amsterdam - www.brainbank.nl) were included in this project.

### Enrichment of protein aggregates

Circulating protein aggregates (CPA) were enriched from plasma using ultracentrifugation (UC) as previously reported (^8^). Briefly, Triton X-100 was added to plasma samples to a final concentration of 2%, then the mixture was incubated for 10 minutes at room temperature and centrifuged at 21000xg for 15 minutes. The supernatant was placed onto a sucrose cushion (1 M sucrose, 50 mM Tris-HCl pH 7.4, 1 mM EDTA and 2% Triton X-100). Ultracentrifugation (UC) was performed for 2 hours at 50000 rpm (167829.2xg) at 4°C, using a Sorvall Discovery 100SE with a TFT 80.2 rotor. The supernatant was discarded, while the pellet was resuspended in PBS (1.5 NaCl) and vortexed for 30 seconds. An additional 40 minutes UC was undertaken for CPA pellet enrichment. The UC final product was resuspended in experimental procedure-specific media including 1) a buffer suitable to investigate aggregates resistance to digestion (detailed below) and 2) the SysQuant Buffer (8M urea, phosphatase inhibitor (PhosSTOP™, Merck) and protease inhibitor (cOmplete™, Merck) to enable proteomic analysis by TMTcalibrator™.

The same protocol was applied to brain samples after mechanical homogenisation in buffer (0.8 M NaCl, 1% Triton X-100, 0.1 M Ethylenediaminetetraacetic acid (EDTA), 0.01 M Tris at pH 7.4 and proteinase inhibitor (cOmplete™, Merck). After mixing buffer and sample in a 10:1 ratio (v/w), the mixture was sonicated at maximum power on ice for 5 minutes and centrifuged at 21000xg for 30 minutes at 4°C. Supernatant was collected and pellet re-suspended in 10 volumes of homogenisation buffer on ice, re-sonicated on ice at maximum power for 30 seconds and finally centrifuged at 21000xg for 30 minutes at 4 °C. Supernatants were finally pooled and used for aggregates enrichment.

#### Quantification of CPA

The protein aggregate fractions resuspended in 8M urea were tested using Pierce™ BCA Protein Assay Kit (ThermoFisher) for total protein quantitation.

#### Transmission Electron Microscopy (TEM)

Qualitative CPA analysis was performed by Transmission Electron Microscopy (TEM). The final UC pellet was resuspended in 500 µl PBS instead of urea. Samples were subjected to an additional washing and UC step (50,000 rpm for 40 minutes, 4°C). The supernatant was then discarded, the pellets re-suspended in 100 µl double distilled water (ddH2O), transferred into a clean tube and sonicated on ice at max power for 5 minutes (Diogenode, Bioruptor) to disrupt possible assembly produced by g-force. Enriched fractions were stored at −80°C for further analysis. A glow-discharged 400 mesh grid coated with carbon was incubated with a droplet of aggregates-enriched sample and after 10 seconds, the excess was removed by carefully touching to the grid edge with filter paper. Negative staining was obtained incubating the grid with a droplet of 2% w/v uranyl acetate (UA). After washing with ddH2O, the grid was air-dried at room temperature and micrographs acquired by a JEOL JEM 1230 electron microscope.

### IgG extraction from plasma and quantification

IgG extraction from the same plasma samples used for CPA separation was carried out using Protein G Spin Columns (Thermo Scientific, UK). Protein G columns were equilibrated by adding 400μL of binding buffer (0.1M phosphate, 0.15M sodium chloride; pH 7.2). After centrifugation, 200μL of plasma diluted 1:2 in PBS was added to the column and incubated at room temperature for 10 minutes. The column was washed three times with binding buffer and the IgG fraction was eluted after addition of 0.1M glycine, pH 2 and collected in neutralization buffer (1M Tris at pH 8-9). The fraction of purified antibody was determined by measuring the relative absorbance of each fraction at 280 nm and the buffer exchanged into PBS using Amicon Ultra centrifugal filter with 100 KDa molecular weight cutoff (Millipore Merck, UK).

### Circulating and brain protein aggregates protease digestion

Aggregates-enriched fractions were enzymatically digested using trypsin (V542A, Promega), α-Chymotrypsin (referred as Chymotrypsin in the text, C4129, Sigma), Calpain (208712, Millipore) and Enterokinase (11334115001, Roche). To minimize UC-induced protease resistance, for disrupting disulphide bonds and enhancing cleavage sites accessibility, pellets were first re-suspended in 50 µl of each protease enzyme recommended buffer (PBS for Trypsin, 100mM Tris HCl for Chymotrypsin, 50mM Hepes - 30mM NaCl for Calpain and 50mM Tris HCl for Enterokinase). 5 µl of 0.5 M DTT was then added prior to sonication. Finally, each enzyme was added into the digestion reaction mix tube at a ratio 1:20 protease:total protein. 5.5 µl of 0.1 M CaCl2 were added to the chymotrypsin and calpain reaction mixes for enzyme activation as indicated by the manufacturers. Digestion mixes were subsequently incubated overnight at 37°C to allow digestion and later stopped adding loading buffer 4X (Fisher Scientific), dithiothreitol (DTT) and by heating at 95°C for 10 minutes. Digested samples were then stored at −80°C for further use. Digestion products were resolved in 3-8% tris-acetate SDS-PAGE followed and visualized by Zinc staining (Life Technologies). Images acquisition (Chemi-Doc Camera, Bio-Rad) and processing (ImageJ) was performed using the Analyze\Gels\Plot lanes function and band intensities obtained with reference to the size marker (HiMark™ Pre-stained Protein Standard). Correction factors to adjust loading differences across samples included the ratio between the sum of all band intensities for each undigested sample and the sum of the marker total band intensities (from the same gel; Supplementary material, Table S5). Resistance to proteases of NfH CPA content was tested by western blotting. Band intensities of NfH digested products were corrected using NfH undigested samples band intensities as reference and normalized for loading volumes differences as reported in the supplementary material (Table S6).

### Western blotting

Proteins loaded onto gels were transferred after electrophoresis to a polyvinylidene difluoride (PVDF) membrane, stained by Ponceau and then blocked with 5% skimmed milk in Tris-Buffered Saline (TBS) 0.1% Tween-20 buffer (TBS-T 0.1%) at room temperature for 1 hour. Overnight incubation was performed with primary antibody at 4 °C followed by incubation with secondary antibody for 1 hour at RT, with membrane washes between steps using TBS-T 0.1%. Lastly, membranes were incubated with enhanced chemiluminescence substrate (ECL). Membrane imaging was undertaken using Image Lab (Bio-Rad) and bands volume measurement using the “Volume Tools” function in Image Lab and “Adj. Vol. (Int)”. Antibodies used in this study are listed in Table S7.

### MS-based proteomics

To evaluate protein composition of the enriched protein aggregate fractions from plasma and brain, we have first undertaken LC-MS/MS analysis after in-gel trypsin digestion of pooled plasma samples (PPS) CPA from of ALS and HC individuals as well as of BPA from ALS individuals. We have then applied TMTcalibrator™ proteomics on individual ALS and HC CPA samples using ALS brain as ion source.

#### In-gel trypsin digestion

Samples were loaded onto a gel for electrophoresis and gel bands were cut out, followed by washing in ammoniumbicarbonate (Ambic): acetonitrile (ACN) and dehydration with ACN. Subsequently, disulfide bonds reduction with DTT and alkylation with Iodoacetamide (IAA) was performed. After washing and complete de-staining, each gel piece was rehydrated with a minimal volume of a solution 50 mM Ambic, 0.01 µg/µl Trypsin (V542A, Promega). Each sample was then incubated overnight (ON) at 37 °C for protein digestion. The following day the tryptic peptides were recovered and each sample was freeze-dried in a vacuum centrifuge for LC-MS/MS analysis as previously described (^8^).

##### Aggregation propensity

To study the chemical properties of the protein mix within the CPA which affect their propensity to aggregate, we have undertaken in-silico analysis of protein size, isoelectric point (pI) and hydrophobicity using Uniprot - ExPASy, which include the Compute pI/MW web tool and GRAVY score (http://www.gravy-calculator.de). This analysis was undertaken comparing the ALS, HC and brain protein lists to the entire Uniprot human proteome (reviewed sequences only).

#### TMTcalibrator™

TMTcalibrator™ workflow was developed by Proteome Sciences plc (^28,29,32^) to quantify low-abundance peptides and proteins in matrices with a high degree of biological complexity. In this experiment, ALS brain tissue was used as calibrant, lysate of two ALS brain samples mixed 1:1, loaded at high concentration along with 6 ALS patients and 6 HC individuals which were considered as analytical samples (Figure S1).

Briefly, the calibrant sample was prepared by dissolving the two brain samples, removing the debris and mixing the two samples 1:1 (w/v):(w/v), while CPA were obtained as described above. All the samples were diluted in SysQuant Buffer in order to process 40µg of total protein for each analytical sample and 840µg for each calibrant sample. Following, reduction with DTT and alkylation with IAA was carried out. After desalting with SepPak tC18, the calibrant volume was divided into four different aliquots with a 1:4:6:10 volume ratio (Calibrant channels), followed by a freeze-dry cycle. Calibrant aliquots contained enough sample for the two 10plexes.

Dried samples were re-solubilised in 120 µl KH2PO4 and TMT reagents were added combining a specific tag for each sample (Figure S1) and incubating at RT for 1 hour. To stop the reactions, hydroxylamine was added in each tube to a final concentration of 0.25% (w/v) and incubated for 15 minutes. At this stage the samples included in each 10plex were merged and were fractionated by basic Reverse Phase (bRP) using Pierce™ High pH Reversed-Phase Peptide Fractionation Kit (ThermoFisher Scientific), generating eight fractions for each of the two 10plexes.

LC-MS/MS analysis was performed in double-shot using a Thermo Scientific™ Orbitrap Fusion Tribrid (Thermo Scientific) mass spectrometer coupled to an EASY-nLC 1000 (Thermo Scientific) system. The 16 bRP fractions were resuspended in 2% ACN, 0.1% formic acid (FA), and then 12 µg from each was injected into a 75 μm × 2 cm nanoViper C18 Acclaim PepMap100 precolumn (3 μm particle size, 100 Å pore size; P/N 164705; Thermo Scientific). Peptides were separated at a flow rate of 250 nl/min and eluted from the column over a 5 hours gradient starting with 0.1% FA in ACN (5-30% ACN from 0 to 280min followed by 10min ramping up to 80% ACN) through a 75 μm × 50 cm PepMap RSLC analytical column at 40 °C (2 μm particle size, 100 Å pore size; P/N ES803; Thermo). After electrospray ionisation, MS spectra ranging from 350 to 1500 m/z values were acquired in the Orbitrap at 120 k resolution and the most intense ions with a minimal required signal of 10,000 were subjected to MS/MS by HCD fragmentation in the Orbitrap at 30 k resolution. Protein identification was carried out with Thermo Scientific Proteome Discoverer 1.4.

##### Bioinformatics

LC-MS/MS analysis generated 32 files that were processed by Proteome Sciences’ proprietary workflows for TMTcalibrator™ including: Calibrator Data Integration Tool (CalDIT), Feature Selection Tool (FeaST) and Functional Analysis Tool (FAT) (^28,32^) (Figure S2). All raw spectra were searched against the human FASTA UniProtKB/Swiss-Prot using SEQUEST-HT and raw intensity values were measured through the TMT reporter ions.

### PC12 viability

Undifferentiated PC12 cells were cultured in Dulbecco’s modified Eagle’s medium (Invitrogen, Paisley, UK) supplemented with 10% fetal calf serum (Invitrogen) and 10% horse serum (Sigma), 100 µg/ml streptomycin, 100 U/ml penicillin (Invitrogen) and incubated at 37° in a 5% CO_2_ humidified atmosphere. These cells were differentiated by plating at a density of 3×10^5^ cells/well in 96-well plates (Nunc, Thermofisher, UK) in Dulbecco’s modified Eagle’s medium (0,1% horse serum supplemented with nerve growth factor (50 ng/ml) and treated for 24 hours at RT with aggregates dissolved in Urea 8M (we assumed that testing cells with aggregates soluble proteins would have been more informative) or with IgG in medium.

Viability after incubation was tested incubating with3-(4,5-dimethylthiazol-2-yl)-2,5diphenyltetrazolium bromide (MTT) at 0.5 mg/ml for 4h, supernatants discarded and 200 µl DMSO added to solubilize the formazan crystals formed after incubation. Colorimetric changes were measured at 590nm (Synergy HT microplate reader). The percentage of cell viability was calculated as the absorbance of treated cells/absorbance of control.

### Endothelial cell viability

Human cerebromicrovascular endothelial cell line hCMEC/D3 was maintained and treated as described previously (^59,60^). Cells were plated on plastic coated with 0.06 µg/cm^2^ calf skin collagen type I (Sigma, UK) and were cultured to confluency in complete endothelial cell growth medium MV2 (PromoCell GmbH, Germany). Following treatment for 24h with aggregates and IgG in medium, cell number was estimated using the Prestoblue HS Cell Viability assay (ThermoFisher Scientific Ltd., UK) according to the manufacturer’s instructions and using a CLARIOstar fluorescence microplate reader (BMG Labtech Ltd., UK) with excitation and emission filters set to 560nm and 590nm respectively.

### Statistics

Proteome data were analysed for normality using the Shapiro-Wilk test. Non-parametric group analysis was performed using Kruskal–Wallis one-way test of variance on ranks with Dunn’s multiple comparison as post-test using GraphPad(v7).

To test ALS versus HC proteolytic bands intensity difference, a “t-Test - Two-Sample Assuming Unequal Variances” was performed.

The enriched KEGG pathways obtained from the submission of ALS and HC pools protein lists to Webgestalt were evaluated for statistical significance using the hypergeometric test (^61^).

Principal component analysis (PCA) was used to study the variance of the data sets generated by the TMTcalibrator™ workflow. To determine the regulated features (FeaST), LIMMA considered the following linear model: logRatio(ALS/HC) ≈ class + group + gender + progression rate + TMT batch. Multiple testing corrections and false discovery rate (FDR) were obtained using the Benjamini-Hochberg procedure. For the Functional analysis (FAT), a two-sided p-value was generated by the Mann Whitney U test and the Benjamini-Hochberg method was used for multiple test correction. Expression values were normalised with other 1000 randomly selected background expression values. A minimum of three matched identifiers (e.g. gene names) were required for each term. Terms with an adjusted p-value < 0.3 were considered significant.

Cell survival assays were evaluated by two-way ANOVA and Tukey HSD test.

### Study approval

A written informed consent was signed by all participants enrolled in the ALS biomarkers study (REC n. 09/H0703/27), including both individuals with a diagnosis of amyotrophic lateral sclerosis (ALS) and healthy controls (HC). Brain had been collected from donors from whom written informed consent for a brain autopsy and the use of the material and clinical information for research purposes had been obtained (NBB: under ethical permission 2009/148).

## Acknowledgments

We would like to thank the patients and their families along with all healthy donors for their contribution. This study is funded by a Medical Research Council (MRC) Industry CASE Studentship (grant number: MR/M015882/1) awarded to Queen Mary University of London and in collaboration with Proteome Sciences. Plasma samples were obtained from the ALS biomarkers study (09/H0703/27). Support for the development of the methodology outlined in this paper has also come from EU2020 funding (H2020 PHC-13-2014): “Efficacy and safety of low-dose IL-2 (ld-IL-2) as a Treg enhancer for anti-neuroinflammatory therapy in newly diagnosed Amyotrophic Lateral Sclerosis (ALS) patients” (MIROCALS)”.

## Supplementary Information

### Study participant: cohort composition

**Table S1.** Clinical and demographic features of the amyotrophic lateral sclerosis (ALS) and healthy controls (HC) individuals selected for the LC-MS proteomic analysis of plasma pool samples. M:F: males (M) and females (F) ratio Diagnostic classification: diagnosis of ALS according to the El-Escorial criteria (^58^) Site of onset: anatomic site of disease onset (e.g. limb vs bulbar). ALS Functional Rating Scale revised: level of neurological impairment across different clinical domains (1-48, higher neurological impairment with lower values) Progression rate at last visit: calculated as 48 - ALSFRS-R score at last visit/disease duration from onset of symptoms to sampling time expressed in months

| Group | M:F | Ethnicity | Age at visit (years) | Diagnostic classification | Site of Onset | ALSFRS-R | Progression rate at visit |
| --- | --- | --- | --- | --- | --- | --- | --- |
| HC | 3:3 | Caucasian (100%) | 58,7 | NA | NA | NA | NA |
| ALS | 5:1 | Caucasian (83,3%), Afro-Caribbean (16,7%) | 65,5 | Definite ALS (33,3%), Possible ALS (33,3%), Probable ALS (16,7%), Suspected ALS (16,7%) | Limb (33,3%), Bulbar (16,7%), Respiratory (16,7%), Bulbar/Respiratory (33,3%) | 39 | 1,974 |

**Table S2.** Clinical and demographic features of the amyotrophic lateral sclerosis (ALS) and healthy controls (HC) individuals selected for CPA digestion and TMTcalibrator™ proteomic analysis. M:F: males (M) and females (F) ratio Diagnostic classification: diagnosis of ALS according to the El-Escorial criteria (^58^) Site of onset: anatomic site of disease onset (e.g. limb vs bulbar). ALS Functional Rating Scale revised: level of neurological impairment across different clinical domains (1-48, higher neurological impairment with lower values) Progression rate at last visit: calculated as 48 - ALSFRS-R score at sampling time at last visit/disease duration from onset of symptoms to sampling time at last visit expressed in months

| Group | M:F | Ethnicity | Age at visit (years) | Diagnostic classification | Site of Onset | ALSFRS-R | Progression rate at visit |
| --- | --- | --- | --- | --- | --- | --- | --- |
| HC | 3:3 | Caucasian (100%) | 64.2 | NA | NA | NA | NA |
| ALS | 3:3 | Caucasian (100%) | 63.8 | Definite ALS (100%) | Limb (100%) | 29 | 0.961 |

**Table S3.** Clinical and demographic features of the amyotrophic lateral sclerosis (ALS) and healthy controls (HC) individuals selected for validation by western blot. M:F: males (M) and females (F) ratio Diagnostic classification: diagnosis of ALS according to the El-Escorial criteria (^58^) Site of onset: anatomic site of disease onset (e.g. limb vs bulbar). ALS Functional Rating Scale revised: level of neurological impairment across different clinical domains (1-48, higher neurological impairment with lower values) Progression rate at last visit: calculated as 48 - ALSFRS-R score at last visit/disease duration from onset of symptoms to last visit expressed in months

| Group | M:F | Ethnicity | Age at visit (years) | Diagnostic classification | Site of Onset | ALSFRS-R | Progression rate at visit |
| --- | --- | --- | --- | --- | --- | --- | --- |
| HC | 2:4 | Caucasian (100%) | 61.8 | NA | NA | NA | NA |
| ALS | 2:2 | Caucasian (100%) | 63.9 | Definite ALS (100%) | Limb (100%) | 27 | 1.082 |

**Table S4.**
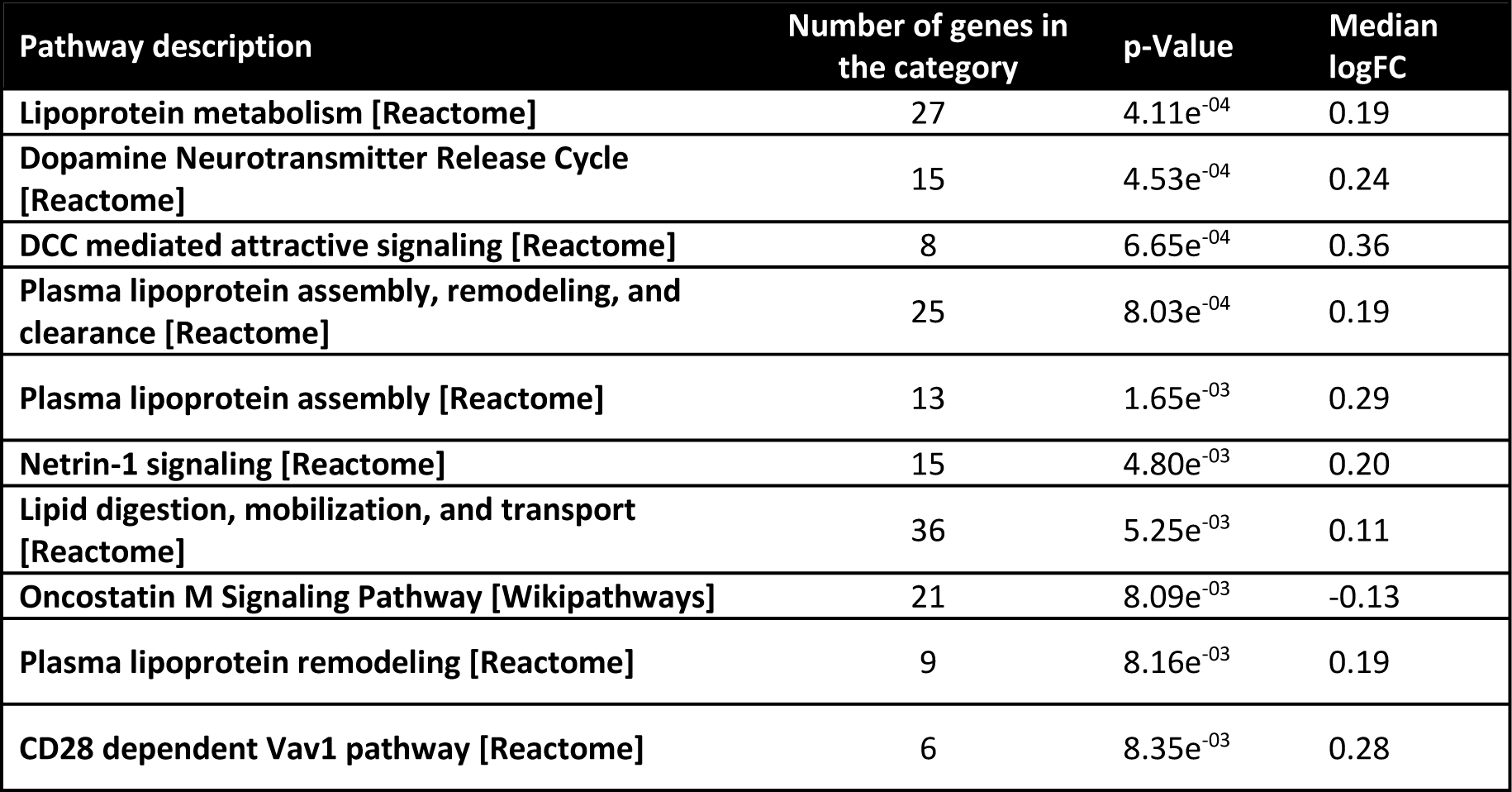
FAT analysis of the ALS vs HC TMT proteomics: top10 regulated pathways. Pathway description: name of the pathway identified (reference database in square brackets, e.g. [Reactome]) Number of genes in the category: genes included in the given pathway in Homo sapiens p-Value: statistical significance calculated by Mann-Whitney U test Median logFC: median value of expression (logFC values) of the proteins included in the given pathway in Homo sapiens

| Pathway description | Number of genes in the category | p-Value | Median logFC |
| --- | --- | --- | --- |
| Lipoprotein metabolism [Reactome] | 27 | 4.11e<sup>-04</sup> | 0.19 |
| Dopamine Neurotransmitter Release Cycle [Reactome] | 15 | 4.53e<sup>-04</sup> | 0.24 |
| DCC mediated attractive signaling [Reactome] | 8 | 6.65e<sup>-04</sup> | 0.36 |
| Plasma lipoprotein assembly, remodeling, and clearance [Reactome] | 25 | 8.03e<sup>-04</sup> | 0.19 |
| Plasma lipoprotein assembly [Reactome] | 13 | 1.65e<sup>-03</sup> | 0.29 |
| Netrin-1 signaling [Reactome] | 15 | 4.80e<sup>-03</sup> | 0.20 |
| Lipid digestion, mobilization, and transport [Reactome] | 36 | 5.25e<sup>-03</sup> | 0.11 |
| Oncostatin M Signaling Pathway [Wikipathways] | 21 | 8.09e<sup>-03</sup> | -0.13 |
| Plasma lipoprotein remodeling [Reactome] | 9 | 8.16e<sup>-03</sup> | 0.19 |
| CD28 dependent Vav1 pathway [Reactome] | 6 | 8.35e<sup>-03</sup> | 0.28 |

### Correction factors for semi-quantitative analysis of neurofilament heavy chain (NfH) in circulating protein aggregates (CPAs) before and after digestion with proteases

For semi-quantitative analysis of ALS and HC digested products, band intensities of the related undigested samples were used as reference to adjust for differences in the total protein content loading across samples (SDS-PAGE was subjected to zinc staining as described in the methods section),. The sum of the intensity of all bands in each lane was compared across samples (Table S5). Excluding the Marker which showed a consistent readout across all gels (CV=6.3%), the loaded undigested CPA samples showed a certain degree of variation (CV>10%, Table S5). To eliminate this variable, a correction factor (CF) was generated as the ratio between the sum of the band intensities acquired from a sample and the sum of the intensities of the Marker in the same gel (Table S6). These factors were applied to the semi-quantitative analysis of NfH expression following CPA digestion. In the late phase of the experiment HC1 and HC2 were eliminated from the analysis as the total protein concentration of these samples was too low. Equal loading for that amount would not allow any NfH detection from any sample by western blot.

**Table S5.** Marker, undigested ALS and HC sample intensities. Total bands intensity is obtained as the sum of the intensities of all bands in a lane (ImageJ) for the gel marker, for ALS and HC undigested samples in each gel (Gel1-6). Mean, standard deviation (St. Dev.) and coefficient of variation (CV%) are shown for each group. The low CV% of the marker confirmed equal loading and the possibility to be used as reliable reference signal. A higher CV% was obtained for ALS and in particular for HC samples, suggesting a slightly more uneven loading. In grey colour-code the Mean, St. Dev. and CV% calculated for the total intensities.

| Total intensity for: | Gel1 (ALS1_HC1) | Gel2 (ALS2_HC2) | Gel3 (ALS3_HC3) | Gel4 (ALS4_HC4) | Gel5 (ALS5_HC5) | Gel6 (HC6) | Mean | St. Dev. | CV% |
| --- | --- | --- | --- | --- | --- | --- | --- | --- | --- |
| Marker | 107704 | 109638 | 102576 | 114133 | 100891 | 95666 | 105102 | 6664 | 6.3 |
| ALS | 62481 | 57628 | 58192 | 55143 | 46977 | - | 56084 | 5735 | 10.2 |
| HC | 40800 | 59041 | 39941 | 46063 | 40622 | 50107 | 46095 | 7477 | 16.2 |

**Table S6.** Correction factors (CFs) for each ALS and HCs sample calculated using undigested samples and marker total intensities as reported in Table S5. The correction factors (CFs) were applied to quantify differences in CPA band intensities of digested samples obtained by SDS-PAGE. Each band under analysis, was first normalized with the marker band intensity closer to the band molecular weight (MW) and then divided by the specific sample/lane CF. ALS1*-5* and HC3-6 indicates the samples used in this experiment with relative CF.

|  | ALS1* | ALS2* | ALS3* | ALS4* | ALS5* | HC3 | HC4 | HC5 | HC6 |
| --- | --- | --- | --- | --- | --- | --- | --- | --- | --- |
| Correction Factors | 0.58 | 0.56 | 0.57 | 0.48 | 0.47 | 0.39 | 0.40 | 0.40 | 0.52 |

### Antibodies used in the study

**Table S7.**
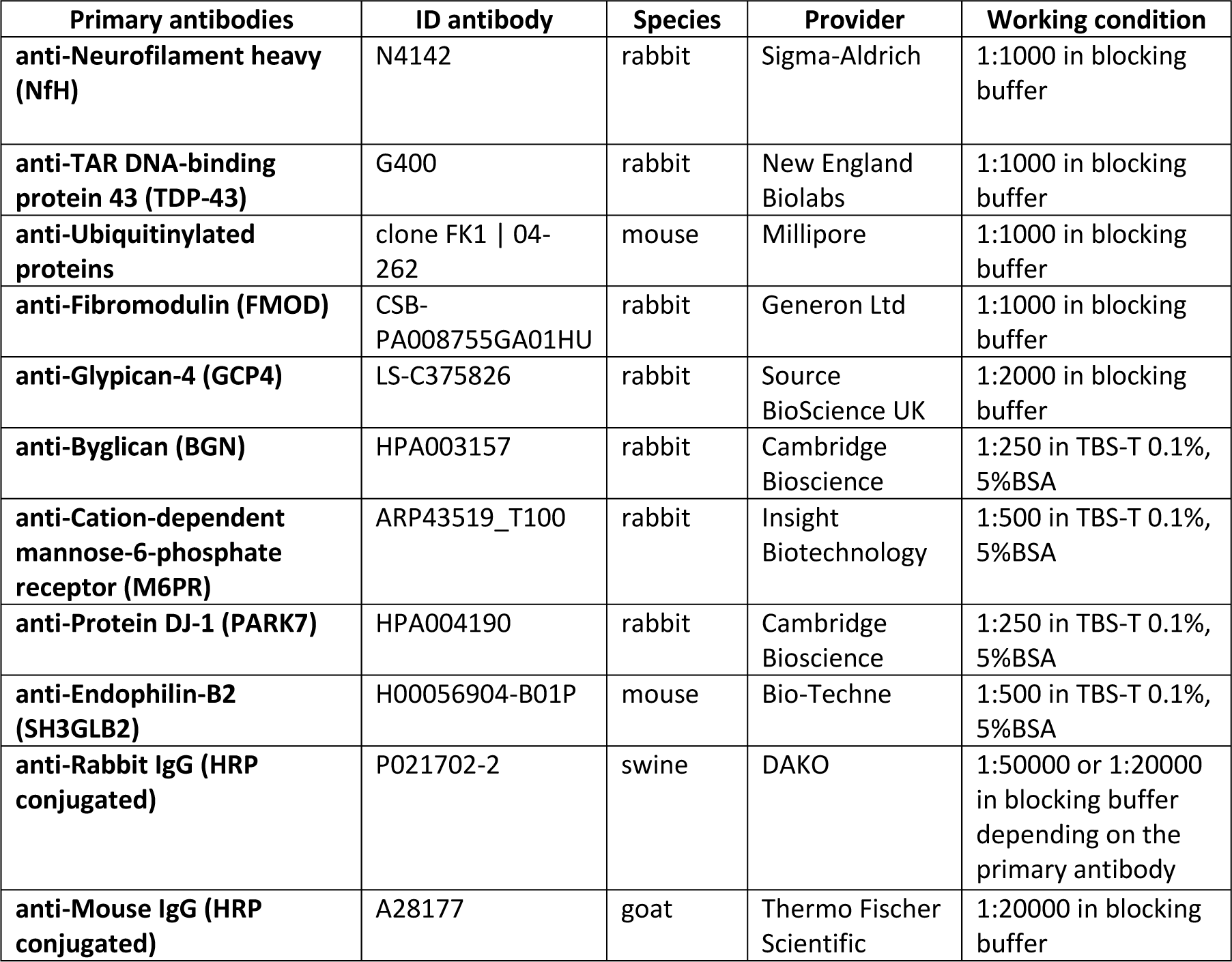
List of antibodies used for western blotting, including primary and secondary antibodies.

## Supplementary figures – TMTcalibrator™

**Figure S1.**
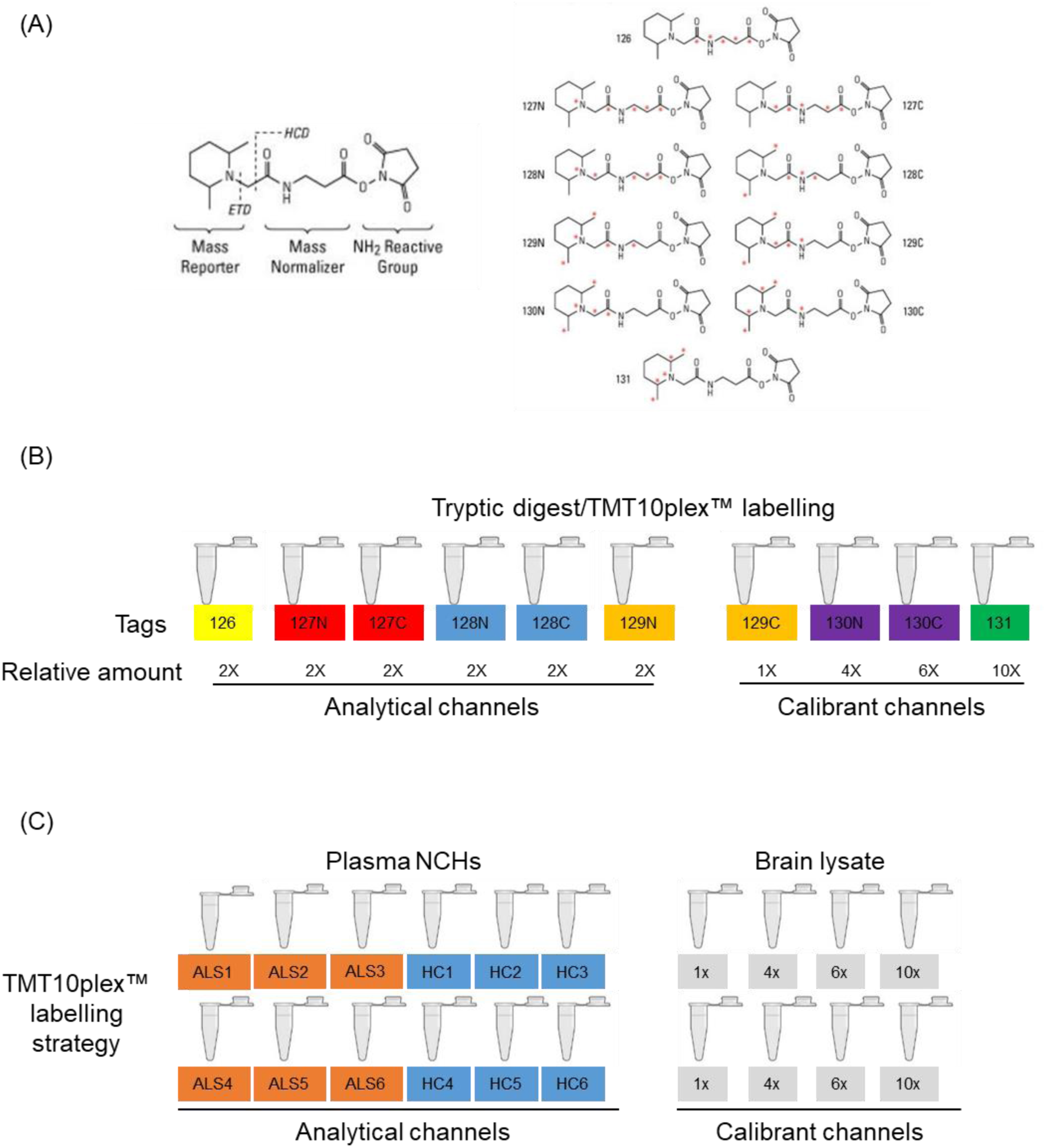
TMTcalibrator™ experimental design. (A) Tandem Mass Tag (TMT) reagents with relative masses and isotope position. (B) General 10plex labelling layout after trypsin digestion of the samples; analytical samples and calilbrants are mixed in a specific ratio that enhances detection by LC-MS/MS of low abundant peptides in the analytical channels thanks to the high calibrant content. (C) Labelling strategy in two 10plexes LC-MS/MS runs which includes Circulating Protein aggregates (CPA) from amyotrophic lateral sclerosis (ALS) patients and from healthy controls (HC) in the analytical channels (orange and blue colour codes) and a mixture (1:1) of brains (Precentral gyrus) lysates from two different ALS patients in the calibrant channels (grey colour code).

**Figure S2.**
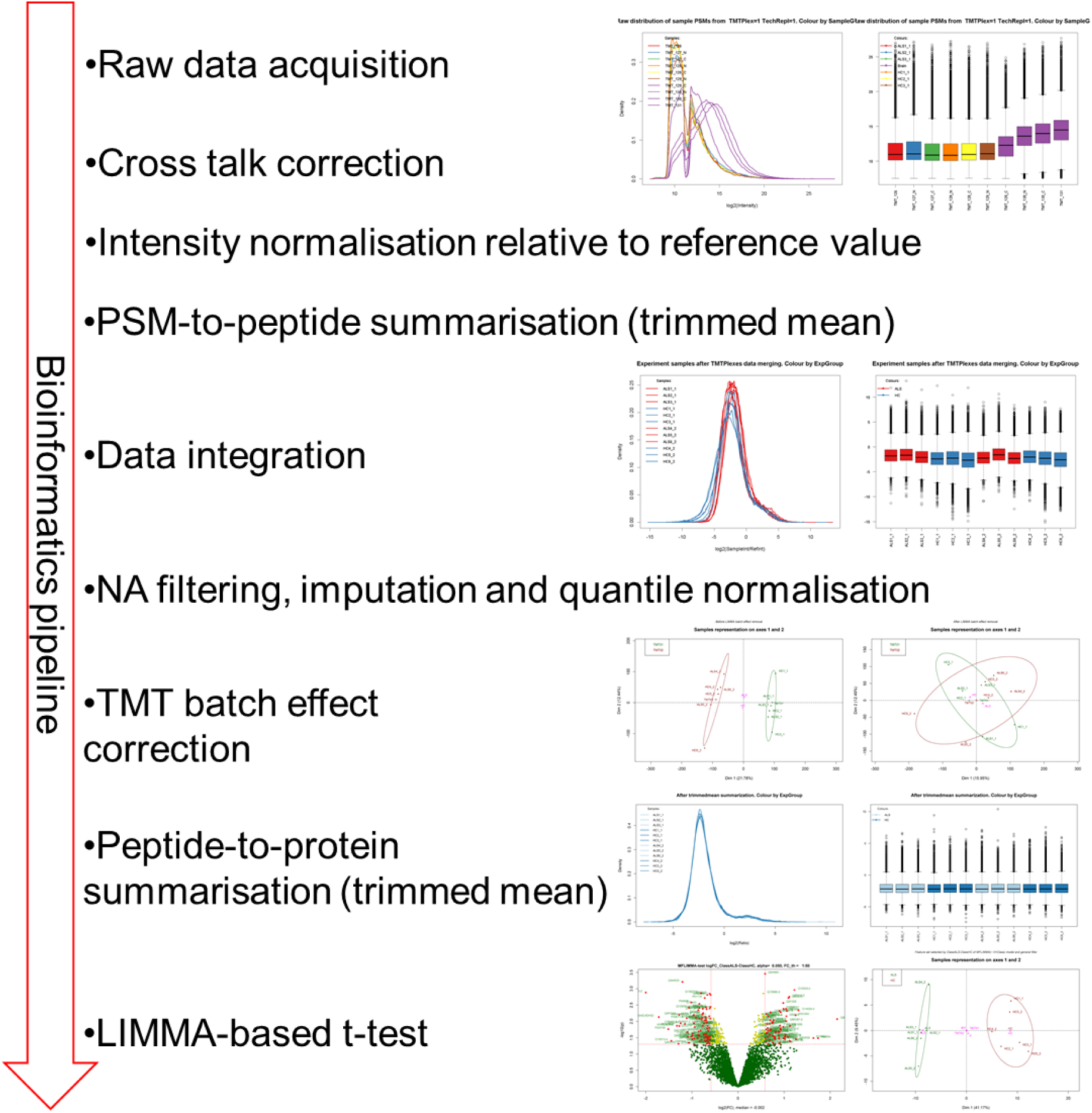
TMTcalibrator™: bioinformatic pipeline. After MS/MS spectra (raw data) acquisition, the intensity of each channel was corrected for background and cross-talking between tags in the second mass spectrometer (MS2). Intensity values of the detected Peptide-Spectrum Matches (PSMs) were normalised with a reference value generated as the average of the Calibrant channels and this was followed by PSM-to-peptide summarisation defined as “trimmed mean”. In this passage the data points considered as outliers in each analytical sample were removed stabilizing the mean before merging the data obtained from the two 10plexes analysed. Then, “not available data points (NA)” filtering, imputation and quantile normalization were performed on the merged data set, so that it was possible to perform a Principal Component Analysis (PCA) on the data acquired. It was also possible to evaluate the TMT batch effect within linear models for microarray data (LIMMA). After peptide-to-protein summarisation, statistical significance of the protein groups identified was analysed with LIMMA-based t-test (p-Value < 0.05, Fold Change threshold = 1.5).

## References

1. Bradley E. Incorporating biomarkers into clinical trial designs: points to consider. Nat Biotechnol. 2012;30(7):596–599. doi:10.1038/nbt.2296

2. Friedrich RP, Tepper K, Rönicke R, et al. Mechanism of amyloid plaque formation suggests an intracellular basis of Abeta pathogenicity. Proc Natl Acad Sci U S A. 2010;107(5):1942–1947. doi:10.1073/pnas.0904532106

3. Lee S, Kim H-J. Prion-like Mechanism in Amyotrophic Lateral Sclerosis: are Protein Aggregates the Key? Exp Neurobiol. 2015;24(1):1–7. doi:10.5607/en.2015.24.1.1

4. Polymenidou M, Cleveland DW. The seeds of neurodegeneration: prion-like spreading in ALS. Cell. 2011;147(3):498–508. doi:10.1016/j.cell.2011.10.011

5. Lu CH, Petzold A, Topping J, et al. Plasma neurofilament heavy chain levels and disease progression in amyotrophic lateral sclerosis: insights from a longitudinal study. J Neurol Neurosurg Psychiatry. 2015;86(5):565–573. doi:10.1136/jnnp-2014-307672

6. Lu CH, Macdonald-Wallis C, Gray E, et al. Neurofilament light chain: A prognostic biomarker in amyotrophic lateral sclerosis. Neurology. 2015;84(22):2247–2257. doi:10.1212/wnl.0000000000001642

7. Yang H, Hu HY. Sequestration of cellular interacting partners by protein aggregates: implication in a loss-of-function pathology. FEBS J. 2016;283(20):3705–3717. doi:10.1111/febs.13722

8. Adiutori R, Aarum J, Zubiri I, et al. The proteome of neurofilament-containing protein aggregates in blood. Biochem Biophys Reports. 2018;14:168–177. doi:10.1016/j.bbrep.2018.04.010

9. Xia K, Trasatti H, Wymer JP, Colon W. Increased levels of hyper-stable protein aggregates in plasma of older adults. Age. 2016;38(3):56. doi:10.1007/s11357-016-9919-9

10. Finn TE, Nunez AC, Sunde M, Easterbrook-Smith SB. Serum albumin prevents protein aggregation and amyloid formation and retains chaperone-like activity in the presence of physiological ligands. J Biol Chem. 2012;287(25):21530–21540. doi:10.1074/jbc.M112.372961

11. Lehallier B, Gate D, Schaum N, et al. Undulating changes in human plasma proteome profiles across the lifespan. Nat Med. 2019;25(12):1843–1850. doi:10.1038/s41591-019-0673-2

12. Williams SA, Kivimaki M, Langenberg C, et al. Plasma protein patterns as comprehensive indicators of health. Nat Med. 2019;25(12):1851–1857. doi:10.1038/s41591-019-0665-2

13. Amor S, Peferoen LAN, Vogel DYS, et al. Inflammation in neurodegenerative diseases - an update. Immunology. 2014. doi:10.1111/imm.12233

14. A. McCombe P, D. Henderson R. The Role of Immune and Inflammatory Mechanisms in ALS. Curr Mol Med. 2011. doi:10.2174/156652411795243450

15. Lyon MS, Wosiski-Kuhn M, Gillespie R, Caress J, Milligan C. Inflammation, Immunity, and amyotrophic lateral sclerosis: I. Etiology and pathology. Muscle and Nerve. 2019. doi:10.1002/mus.26289

16. Villeda SA, Luo J, Mosher KI, et al. The ageing systemic milieu negatively regulates neurogenesis and cognitive function. Nature. 2011. doi:10.1038/nature10357

17. Conboy IM, Conboy MJ, Wagers AJ, Girma ER, Weismann IL, Rando TA. Rejuvenation of aged progenitor cells by exposure to a young systemic environment. Nature. 2005. doi:10.1038/nature03260

18. Middeldorp J, Lehallier B, Villeda SA, et al. Preclinical assessment of young blood plasma for Alzheimer disease. JAMA Neurol. 2016. doi:10.1001/jamaneurol.2016.3185

19. Garbuzova-Davis S, Hernandez-Ontiveros DG, Rodrigues MC, et al. Impaired blood-brain/spinal cord barrier in ALS patients. Brain Res. 2012;1469:114–128. doi:10.1016/j.brainres.2012.05.056

20. Terry C, Wenborn A, Gros N, et al. Ex vivo mammalian prions are formed of paired double helical prion protein fibrils. Open Biol. 2016. doi:10.1098/rsob.160035

21. Safar JG, Wille H, Geschwind MD, et al. Human prions and plasma lipoproteins. Proc Natl Acad Sci. 2006. doi:10.1073/pnas.0604021103

22. Quintana C, Cowley JM, Marhic C. Electron nanodiffraction and high-resolution electron microscopy studies of the structure and composition of physiological and pathological ferritin. J Struct Biol. 2004. doi:10.1016/j.jsb.2004.03.001

23. Sana B, Poh CL, Lim S. A manganese-ferritin nanocomposite as an ultrasensitive T2contrast agent. Chem Commun. 2012. doi:10.1039/c1cc15189d

24. Zhou Z, Fan J-B, Zhu H-L, et al. Crowded Cell-like Environment Accelerates the Nucleation Step of Amyloidogenic Protein Misfolding. J Biol Chem. 2009;284(44):30148–30158. doi:10.1074/jbc.M109.002832

25. Weids AJ, Ibstedt S, Tamas MJ, Grant CM. Distinct stress conditions result in aggregation of proteins with similar properties. Sci Rep. 2016;6:24554. doi:10.1038/srep24554

26. McKinley MP, Bolton DC, Prusiner SB. A protease-resistant protein is a structural component of the scrapie prion. Cell. 1983;35(1):57–62.

27. Lu CH, Kalmar B, Malaspina A, Greensmith L, Petzold A. A method to solubilise protein aggregates for immunoassay quantification which overcomes the neurofilament “hook” effect. J Neurosci Methods. 2011;195(2):143–150. doi:10.1016/j.jneumeth.2010.11.026

28. Leoni E, Bremang M, Mitra V, et al. Combined Tissue-Fluid Proteomics to Unravel Phenotypic Variability in Amyotrophic Lateral Sclerosis. Sci Rep. 2019;9(1). doi:10.1038/s41598-019-40632-4

29. Russell CL, Mitra V, Hansson K, et al. Comprehensive Quantitative Profiling of Tau and Phosphorylated Tau Peptides in Cerebrospinal Fluid by Mass Spectrometry Provides New Biomarker Candidates. Iqbal K, ed. J Alzheimer’s Dis. 2016;55(1):303–313. doi:10.3233/JAD-160633

30. Zubiri I, Lombardi V, Bremang M, et al. Tissue-enhanced plasma proteomic analysis for disease stratification in amyotrophic lateral sclerosis. Mol Neurodegener. 2018;13(1). doi:10.1186/s13024-018-0292-2

31. Database MT human disease. ALS elite genes. https://www.malacards.org/card/amyotrophic_lateral_sclerosis_1#RelatedGenes-table.

32. Zubiri I, Lombardi V, Bremang M, et al. Tissue-enhanced plasma proteomic analysis for disease stratification in amyotrophic lateral sclerosis. 2018;2:1–17.

33. Szelechowski M, Amoedo N, Obre E, et al. Metabolic Reprogramming in Amyotrophic Lateral Sclerosis. Sci Rep. 2018. doi:10.1038/s41598-018-22318-5

34. Tefera TW, Borges K. Metabolic dysfunctions in amyotrophic lateral sclerosis pathogenesis and potential metabolic treatments. Front Neurosci. 2017. doi:10.3389/fnins.2016.00611

35. Delaye JB, Patin F, Piver E, et al. Low IDL-B and high LDL-1 subfraction levels in serum of ALS patients. J Neurol Sci. 2017. doi:10.1016/j.jns.2017.07.019

36. DeWitt DA, Richey PL, Praprotnik D, Silver J, Perry G. Chondroitin sulfate proteoglycans are a common component of neuronal inclusions and astrocytic reaction in neurodegenerative diseases. Brain Res. 1994;656(1):205–209. doi:10.1016/0006-8993(94)91386-2

37. Ancsin JB. Amyloidogenesis: Historical and modern observations point to heparan sulfate proteoglycans as a major culprit. Amyloid. 2003;10(2):67–79. doi:10.3109/13506120309041728

38. Sarrazin S, Lamanna WC, Esko JD. Heparan sulfate proteoglycans. Cold Spring Harb Perspect Biol. 2011;3(7):1–33. doi:10.1101/cshperspect.a004952

39. Hirano K, Ohgomori T, Kobayashi K, et al. Ablation of Keratan Sulfate Accelerates Early Phase Pathogenesis of ALS. PLoS One. 2013. doi:10.1371/journal.pone.0066969

40. Holmes BB, DeVos SL, Kfoury N, et al. Heparan sulfate proteoglycans mediate internalization and propagation of specific proteopathic seeds. Proc Natl Acad Sci. 2013. doi:10.1073/pnas.1301440110

41. Forostyak S, Homola A, Turnovcova K, Svitil P, Jendelova P, Sykova E. Intrathecal delivery of mesenchymal stromal cells protects the structure of altered perineuronal nets in SOD1 rats and amends the course of ALS. Stem Cells. 2014. doi:10.1002/stem.1812

42. Foyez T, Takeda-Uchimura Y, Ishigaki S, et al. Microglial keratan sulfate epitope elicits in central nervous tissues of transgenic model mice and patients with amyotrophic lateral sclerosis. Am J Pathol. 2015;185(11):3053–3065. doi:10.1016/j.ajpath.2015.07.016

43. Nishitsuji K. Heparan sulfate S-domains and extracellular sulfatases (Sulfs): their possible roles in protein aggregation diseases. Glycoconj J. 2018:387–396. doi:10.1007/s10719-018-9833-8

44. Shijo T, Warita H, Suzuki N, et al. Aberrant astrocytic expression of chondroitin sulfate proteoglycan receptors in a rat model of amyotrophic lateral sclerosis. J Neurosci Res. 2018;96(2):222–233. doi:10.1002/jnr.24127

45. Sasaki S. Autophagy in spinal cord motor neurons in sporadic amyotrophic lateral sclerosis. J Neuropathol Exp Neurol. 2011;70. doi:10.1097/NEN.0b013e3182160690

46. Song C, Guo J, Liu Y, Tang B. Autophagy and Its Comprehensive Impact on ALS. Int J Neurosci. 2012;122(12):695–703. doi:10.3109/00207454.2012.714430

47. Cipolat Mis MS, Brajkovic S, Frattini E, Di Fonzo A, Corti S. Autophagy in motor neuron disease: Key pathogenetic mechanisms and therapeutic targets. Mol Cell Neurosci. 2016;72:84–90. doi:10.1016/J.MCN.2016.01.012

48. Sullivan PM, Zhou X, Robins AM, et al. The ALS/FTLD associated protein C9orf72 associates with SMCR8 and WDR41 to regulate the autophagy-lysosome pathway. Acta Neuropathol Commun. 2016. doi:10.1186/s40478-016-0324-5

49. Shi Y, Lin S, Staats KA, et al. Haploinsufficiency leads to neurodegeneration in C9ORF72 ALS/FTD human induced motor neurons. Nat Med. 2018. doi:10.1038/nm.4490

50. Blasco H, Veyrat-Durebex C, Bocca C, et al. Lipidomics Reveals Cerebrospinal-Fluid Signatures of ALS. Sci Rep. 2017. doi:10.1038/s41598-017-17389-9

51. Supattapone S. Phosphatidylethanolamine as a prion cofactor: Potential implications for disease pathogenesis. Prion. 2012. doi:10.4161/pri.21826

52. Saez I, Vilchez D. The Mechanistic Links Between Proteasome Activity, Aging and Age-related Diseases. Curr Genomics. 2014;15(1):38–51. doi:10.2174/138920291501140306113344

53. Ling SC, Polymenidou M, Cleveland DW. Converging mechanisms in ALS and FTD: disrupted RNA and protein homeostasis. Neuron. 2013;79(3):416–438. doi:10.1016/j.neuron.2013.07.033

54. Ngo ST, Steyn FJ. The interplay between metabolic homeostasis and neurodegeneration: insights into the neurometabolic nature of amyotrophic lateral sclerosis. Cell Regen. 2015;4(1):5. doi:10.1186/s13619-015-0019-6

55. Palamiuc L, Schlagowski A, Ngo ST, et al. A metabolic switch toward lipid use in glycolytic muscle is an early pathologic event in a mouse model of amyotrophic lateral sclerosis. EMBO Mol Med. 2015;7(5):526–546. doi:10.15252/emmm.201404433

56. Lu C-H, Petzold A, Kalmar B, Dick J, Malaspina A, Greensmith L. Plasma Neurofilament Heavy Chain Levels Correlate to Markers of Late Stage Disease Progression and Treatment Response in SOD1(G93A) Mice that Model ALS. PLoS One. 2012;7(7):e40998. doi:10.1371/journal.pone.0040998

57. Petzold A, Tisdall MM, Girbes AR, et al. In vivo monitoring of neuronal loss in traumatic brain injury: A microdialysis study. Brain. 2011. doi:10.1093/brain/awq360

58. Ludolph A, Drory V, Hardiman O, et al. A revision of the El Escorial criteria - 2015. Amyotroph Lateral Scler Frontotemporal Degener. 2015;16(5-6):291–292. doi:10.3109/21678421.2015.1049183

59. Weksler BB, Subileau EA, Perrière N, et al. Blood-brain barrier-specific properties of a human adult brain endothelial cell line. FASEB J. 2005;19(13):1872–1874. doi:10.1096/fj.04-3458fje

60. Hoyles L, Snelling T, Umlai U-K, et al. Microbiome–host systems interactions: protective effects of propionate upon the blood–brain barrier. Microbiome 2018 61. 2018;6(1):55. doi:10.1186/s40168-018-0439-y

61. Zhang B, Kirov S, Snoddy J. WebGestalt: An integrated system for exploring gene sets in various biological contexts. Nucleic Acids Res. 2005;33(SUPPL. 2):741–748. doi:10.1093/nar/gki475

